# Origins and evolutionary trajectories of morbilliviruses in Neotropical bats

**DOI:** 10.1101/2025.03.18.643905

**Authors:** Wendy K. Jo, Andres Moreira-Soto, Angélica Cristine Almeida Campos, Luiz Gustavo Bentim Góes, Maria Angélica Mares-Guia, Andrea Rasche, Ana Maria Bispo de Filippis, Gabriela Hernández-Mora, Sham Nambulli, Daniel G. Streicker, W. Paul Duprex, Jan Felix Drexler, bat morbillivirus consortium

## Abstract

Bats are important reservoirs of paramyxoviruses, yet their role in the evolutionary origins of morbilliviruses like measles and rinderpest viruses remains unclear. Investigating wild bats (38/1,629 RT-PCR positive) and non-human primates (NHP, 13/1,370 RT-PCR-positive) from Latin America coupled with data mining, we identified six divergent morbilliviruses. High morbillivirus concentrations in bat organs and morbillivirus RNA staining in NHP livers suggested systemic infection. Of 117 vampire bats, 35.9% had neutralizing antibodies against a primary vampire bat morbillivirus isolate, suggesting frequent survived infections. Usage of bat, but not human, CD150 for cell entry and partial cross-neutralization of bat-associated morbillivirus by heterologous sera suggested conserved entry and antigenicity. Macro-evolutionary reconstructions revealed a predominant role of Neotropical bats during morbilliviral diversification, including bat-associated host shifts into Mexican pigs and Brazilian NHPs. These data argue for increased surveillance, experimental risk assessments, vaccine development, and intervention strategies to cease future outbreaks of previously reservoir-bound morbilliviruses.

## Introduction

Morbilliviruses are highly contagious, mammalian pathogens, including measles virus (MV), rinderpest virus (RPV), peste des petits ruminants virus (PPRV), and canine distemper virus (CDV), which can cause large outbreaks, some of which with high mortality rates^1^. Live-attenuated vaccines are available for the diseases caused by those morbilliviruses and enabled RPV eradication by 2011^2^, whereas MV eradication has not been accomplished yet^3^. MV causes >100,000 human deaths per year and significant morbidity^4^. PPRV is estimated to lead to economic loses between 1.2-1.7 billion USD^5^. RPV caused large epizootics in cattle which led to famines associated with millions of human deaths^2^. CDV infects dogs globally and causes epizootics including cross-species transmission among wild and domestic carnivores^6^. Other morbilliviruses include phocine distemper virus (PDV), cetacean morbillivirus (CeMV), feline morbillivirus (FeMV)^7^ and the recently described porcine morbillivirus (PoMV) found in 2020 during an investigation of fetal damage, encephalitis and stillbirth in a pig breeding facility in northern Mexico^8^. Most recently, morbilliviruses were discovered in shrews and Neotropical bats (**Supplementary Table S1**)^9–13^, but little is known of their evolutionary history or epidemiology.

Bats are important reservoirs of zoonotic paramyxoviruses such as henipaviruses^10^. Bats belong to a mammalian order within the clade Laurasiatheria that radiated about 60 million years ago^14^. They are physiologically and ecologically distinct among mammals and thus their tolerance to some viral infections may be altered^15^. All known bat morbilliviruses are from the Neotropics^9–11^, and their role in the origins of morbilliviruses is unclear. Here, we explored the significance of Neotropical bats in the diversification and origin of morbilliviruses by combining field surveys, data mining, serologic testing, *in vitro* investigations, and evolutionary analyses. We show that Neotropical bats are predominant sources of cross-order host shifts defining the genealogy of morbilliviruses. Our data argue for considerations of future host shifts of previously reservoir-bound morbilliviruses within eradication concepts.

## Results

### Identification of bat-associated morbilliviruses

We investigated 1,629 wild bats representing 16 species sampled for virus surveillance in Brazil and Costa Rica over 14 years (**Fig. 1a**; **Table 1**). The overall RT-PCR-based^10^ detection rate of morbilliviruses was 2.3 % (38/1,629; 95% CI 1.7-3.2). Four bat species belonging to three genera tested positive, including vampire (*Desmodus rotundus*) (4.3%; 33/777, 95% CI 3.0-5.9), mouse-eared (*Myotis nigricans*) (8.8%; 3/34; 95% CI 3.1-22.9), and black mastiff bats (*Molossus rufus*) (1.2%; 2/173; 95% CI 0.32-4.1) (**Fig. 1a**; **Supplementary Table S2**). High virus concentrations of up to 10^9^ RNA copies/g were found in the spleen, the most frequently organ positive by RT-PCR (**Fig. 1b**; **Supplementary Table S2**). Multiple organs including kidney, lung, liver, intestine, and heart were positive at comparable concentrations, suggesting systemic infection similar to other morbilliviruses^16^. Available sera from morbillivirus-positive animals (n=7) were mostly RT-PCR negative or had low virus concentrations, close to the limit of detection. This can be explained by the dissemination of morbilliviruses mainly by lymphocytes^16^ which are absent from sera.

**Fig. 1.**
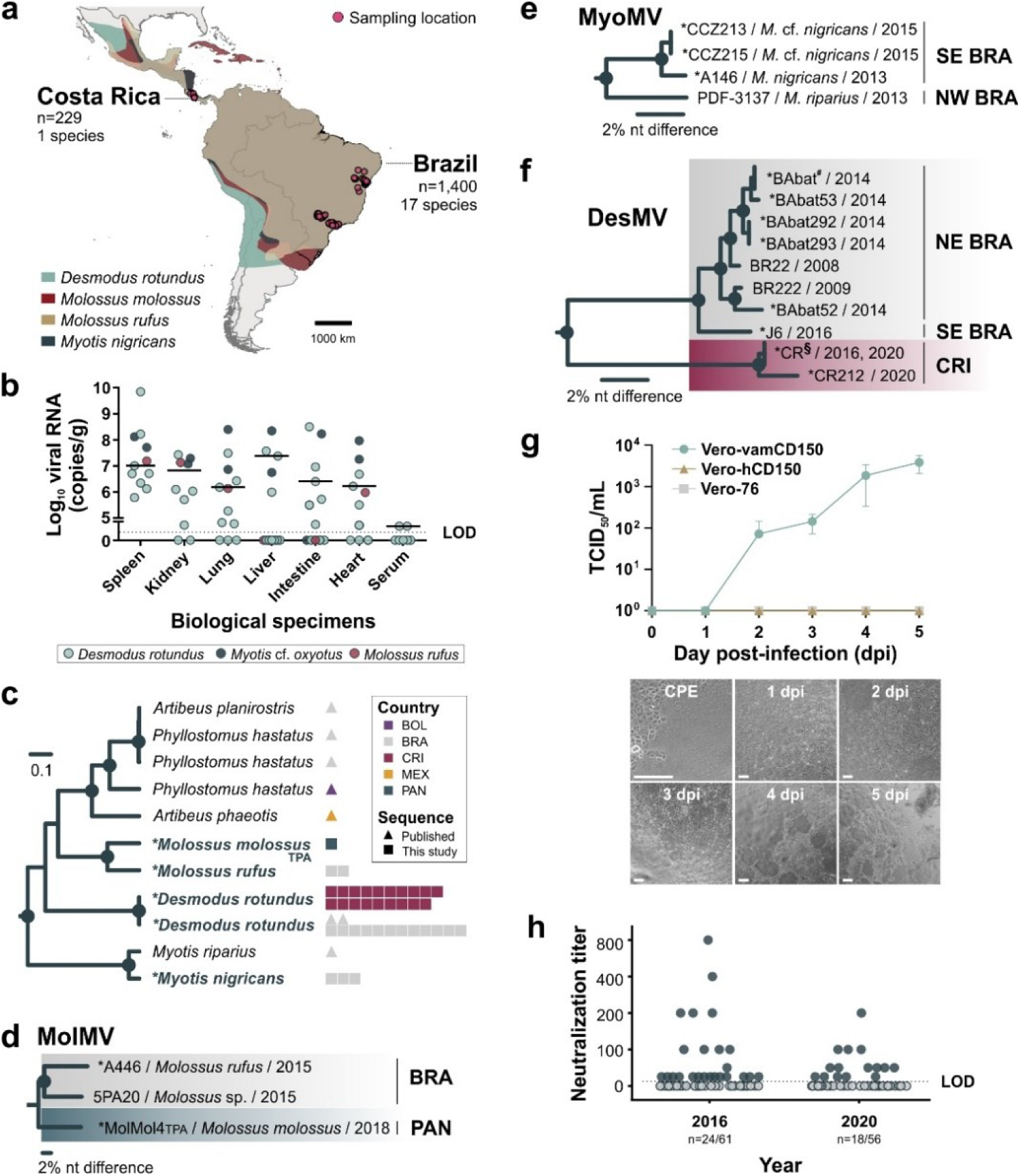
Bat-associated morbillivirus epidemiology. a) Geographical distribution of bat host species of morbilliviruses, sampling countries, and total number of bat species and individuals investigated. b) Viral loads of bat morbilliviruses in different organs from individual bats (n=12). Black line, median. LOD, limit of detection. c) Bayesian phylogeny of bat morbillivirus previously published (triangle) and own data (square). Scale bar indicates genetic distance. TPA, third party annotation. d) Phylogenetic tree of morbillivirus-positive *Molossus* bats from Brazil (BRA) and Panama (PAN) e) Phylogenetic tree of morbillivirus-positive *Myotis* bats from northwestern (NW) and southeastern (SE) BRA. f) Phylogenetic tree of morbillivirus-positive vampire bats (*D. rotundus*) from northeastern (NE) and SE BRA and Costa Rica (CRI). Collapsed taxa at node with symbols: # = BAbat447, -443, -446, -445, -408, -153, -444, - 438, -222, -218; and § = CR123, -128, -163, -183, -191, -205, -207, -209, -217, -220, -221. For panels c-f, partial *L* gene sequences were used and asterisk (*) describes sequences from this study. For panels d-f, Neighbor-joining method was used and circles at nodes indicate bootstrap values ≥80%. See **Supplementary Tables S1-S2** for virus abbreviations and GenBank accession nos. g) Multistep growth curve analyses of DesMV inoculated at a moi of 0.01 for five days in Vero-vamCD150, Vero-hCD150 and Vero-76 as control. Error bars are in standard error of the mean. Representative images of cytopathic effects (syncytia formation) at each day post-infection. Scale bars, 200µm. h) Neutralization titers against DesMV in Costa Rican vampire bat sera. Experiment was performed in duplicate.

**Table 1.** Bat samples.

| Species | Country | States/Province | Total | Pos | Years (n) | Study |
| --- | --- | --- | --- | --- | --- | --- |
| *Artibeus lituratus* | Brazil | Sao Paulo | 31 |  | 2007 (3), 2010 (1), 2013 (3), 2014 (9), 2015 (15) | This study, <sup>67-69</sup> |
| *Artibeus planirostris* | Brazil | Sao Paulo | 5 |  | 2011 (2), 2012 (1), 2013 (1), 2014 (1) | This study, <sup>68,69</sup> |
| *Carollia brevicauda* | Brazil | Bahia | 39 |  | 2009 | <sup>70</sup> |
| *Carollia perspicillata* | Brazil | Bahia | 75 |  | 2009 | <sup>70</sup> |
|  |  | Sao Paulo | 20 |  | 2007 (17), 2011 (1), 2015 (2) | This study, <sup>67-69</sup> |
| *Cynomops planirostris* | Brazil | Sao Paulo | 11 |  | 2013 (1), 2014 (7), 2015 (3) | This study, <sup>69</sup> |
| *Desmodus rotundus* | Brazil | Bahia | 481 | 18 | 2008 (15), 2009 (13), 2010 (2), 2012 (451) | This study, <sup>70</sup> |
|  |  | Sao Paulo | 67 | 1 | 2007 (3), 2011 (41), 2014 (15), 2015 (8) | This study, <sup>67-69,71</sup> |
|  | Costa Rica | Alajuela | 35 |  | 2016 | This study |
|  |  | Cartago | 62 |  | 2016 | This study |
|  |  | Puntarenas | 132 | 14 | 2016 (61), 2020 (71) | This study |
| *Eptesicus furinalis* | Brazil | Sao Paulo | 13 |  | 2011 (2), 2013 (6), 2014 (2), 2015 (3) | This study, <sup>68,69</sup> |
| *Eumops glaucinus* | Brazil | Sao Paulo | 104 |  | 2009 (1), 2010 (1), 2011 (5), 2012 (5), 2013 (19), 2014 (34), 2015 (39) | This study, <sup>67-69</sup> |
| *Eumops perotis* | Brazil | Sao Paulo | 12 |  | 2013 (1), 2014 (8), 2015 (3) | This study, <sup>67,69</sup> |
| *Glossophaga soricina* | Brazil | Bahia | 1 |  | 2009 | <sup>70</sup> |
|  |  | Sao Paulo | 63 |  | 2007 (1), 2011 (2), 2012 (1), 2013 (2), 2014 (26), 2015 (31) | This study, <sup>67-69</sup> |
| *Molossus currentium* | Brazil | Bahia | 5 |  | 2009 | <sup>70</sup> |
| *Molossus molossus* | Brazil | Sao Paulo | 232 |  | 2010 (1), 2011 (24), 2012 (15), 2013 (60), 2014 (80), 2015 (52) | This study, <sup>67-69</sup> |
| *Molossus rufus* | Brazil | Bahia | 17 |  | 2009 | <sup>70</sup> |
|  |  | Sao Paulo | 156 | 2 | 2009 (10), 2010 (1), 2011 (19), 2012 (27), 2013 (48), 2014 (27), 2015 (24) | This study, <sup>67-69</sup> |
| *Myotis nigricans* | Brazil | Sao Paulo | 34 | 3 | 2012 (4), 2013 (10), 2014 (5), 2015 (15) | This study, <sup>68,69</sup> |
| *Platyrrhinus lineatus* | Brazil | Sao Paulo | 5 |  | 2014 (4), 2015 (1) | This study, <sup>68,69</sup> |
| *Tadarida brasiliensis* | Brazil | Sao Paulo | 29 |  | 2014 (14), 2015 (15) | This study, <sup>69</sup> |
| **Total** |  |  | **1,629** | **38** |  |  |

Pos, Morbillivirus positive animals

To complement our field and laboratory investigations, we performed a parallel search for novel, unreported morbilliviruses using Serratus Explorer and palmID in the Serratus platform^17^. The platform uses genomic sequencing data from the National Center for Biotechnology Information Short Read Archive (SRA) and screens for sequences with similarity to known viruses. A divergent morbillivirus was uncovered in sequences derived by RNA-Seq of the blood of a Pallas’s mastiff bat (*Molossus molossus*, SRR11526513) from Panama sampled in 2018. Beyond sequences corresponding to MV, PPRV, and CDV in prototypic hosts, no other divergent morbillivirus sequence was identified in any other animal hosts, including 3,906 SRA accessions from 181 bat species of which 81 (2,004 accessions) were from the Old World and 100 (1,902 accessions) from the Neotropics.

In sum, we detected five divergent bat-associated morbilliviruses, four from own fieldwork, and one by data mining, almost doubling the known morbillivirus bat host diversity and expanding the number of individual bats 4-fold, enabling robust epidemiological, pathogenetic and macro-evolutionary analyses (**Fig. 1c**). All morbillivirus-positive bat species were from the Neotropics, thus highlighting the importance of bats from this area as morbillivirus hosts.

### Epidemiology of bat-associated morbilliviruses

Morbillivirus maintenance in bats is suggested by the detection of genetically closely related *Molossus*- and *Myotis*-associated morbilliviruses (MolMV, MyoMV) in at least two *Molossus* and three different *Myotis* species (**Fig. 1d-e**), reminiscent of bat coronaviruses also capable of infecting multiple species within a host genus^18^. *Desmodus*-associated morbilliviruses (DesMV) were detected across a period spanning 13 years (2008-2020) and grouped in two country-associated clades that shared a most recent common ancestor. The consistent detection and wide geographic extent of DesMV, spanning approximately 6,500 km longitude from Costa Rica over Brazil, supports the possibility of morbillivirus maintenance across the entire range of this species (**Fig. 1f**).

The isolation of viruses from reservoir hosts is challenging due to lack of established tools and specimen degradation. We successfully isolated DesMV from a vampire bat lung in Vero cells expressing the vampire bat CD150 orthologue, showing canonical CD150-mediated cell entry. No utilization of human CD150 by the DesMV isolate was shown (**Fig. 1g**). Substantial degradation of other bat morbillivirus-positive specimens due to sampling under tropical conditions and repeated freeze-thawing cycles prevented virus isolation. We detected the presence of neutralizing antibodies against DesMV in 35.9% (42/117, 95% CI 27.8-44.9) of vampire bats from Costa Rica (**Fig. 1h**), suggesting both frequent exposure and survival of past bat morbillivirus infections. The antibody detection rate in vampire bats was within the seroprevalence ranges detected for other morbilliviruses in wildlife populations^19^. Densely populated roosts and usage of multiple roosts may facilitate virus transmission within a bat colony^20^ and in sympatric bat species through droplets or aerosol containing viruses shed in urine or feces^21^.

### Prototypic genomic properties are conserved across divergent bat-associated morbilliviruses

We obtained complete and near-complete genomes of five bat-associated morbilliviruses by RT-PCR, high throughput sequencing, and SRA data mining. According to current taxonomic species demarcation criteria^1^, branch lengths of >0.03 in the L protein-based phylogeny indicated the overall presence of six morbillivirus species in bats at the current knowledge (**Extended Data** Fig. 1a), namely *Desmodus morbillivirus 1* and *2*, *Molossus morbillivirus 1 and 2*, *Myotis morbillivirus*^9^, and *Phyllostomus morbillivirus*^9^ including our and recently published data^9^ . Based on translated, concatenated ORFs, bat-associated morbilliviruses showed pairwise sequence distances of >30% from most morbilliviruses (**Extended Data Table 1**), comparable to the distance between most other morbillivirus species. DesMV-1 and DesMV-2, which were identified in the same host species but in two geographic regions, showed pairwise amino acid sequence distance of about 11% (**Extended Data Table 1**), suggesting diversification might have occurred within this host species. The number of morbilliviral species in bats exceeded those known in any other host order, i.e., six compared to four in artiodactyls (RPV, PPRV, CeMV, PoMV), three in carnivores (CDV, PDV, and FeMV) and one in primates (MV). The bat-associated morbilliviruses showed characteristic morbillivirus genome organization^22^, with genome lengths of about 16 kb encoding the six structural nucleocapsid (N), phospho-(P), matrix (M), fusion (F), hemagglutinin (H), and L proteins, as well as non-structural proteins (C and V) (**Fig. 2a**). One of the previously published MyoMV sequences (termed PREDICT/PDF-3137) had an extended amino-terminus of the F glycoprotein open reading frame (ORF), like CDV and PDV^22^. In contrast, the genetically related MyoMV from this study and other bat morbilliviruses did not show amino-terminal F glycoprotein extensions. Like all morbilliviruses except for FeMV^7^, bat-associated morbilliviruses contained a polybasic furin cleavage signal in their F glycoprotein (**Extended Data Fig. 1b**). Bat-associated morbilliviruses showed highest divergence in the regions within the highly disordered carboxyl-terminal domain of the N protein and amino-terminal domain of the P protein^23^, and the receptor binding H glycoprotein, potentially associated with host adaptation (**Fig. 2b**).

**Fig. 2.**
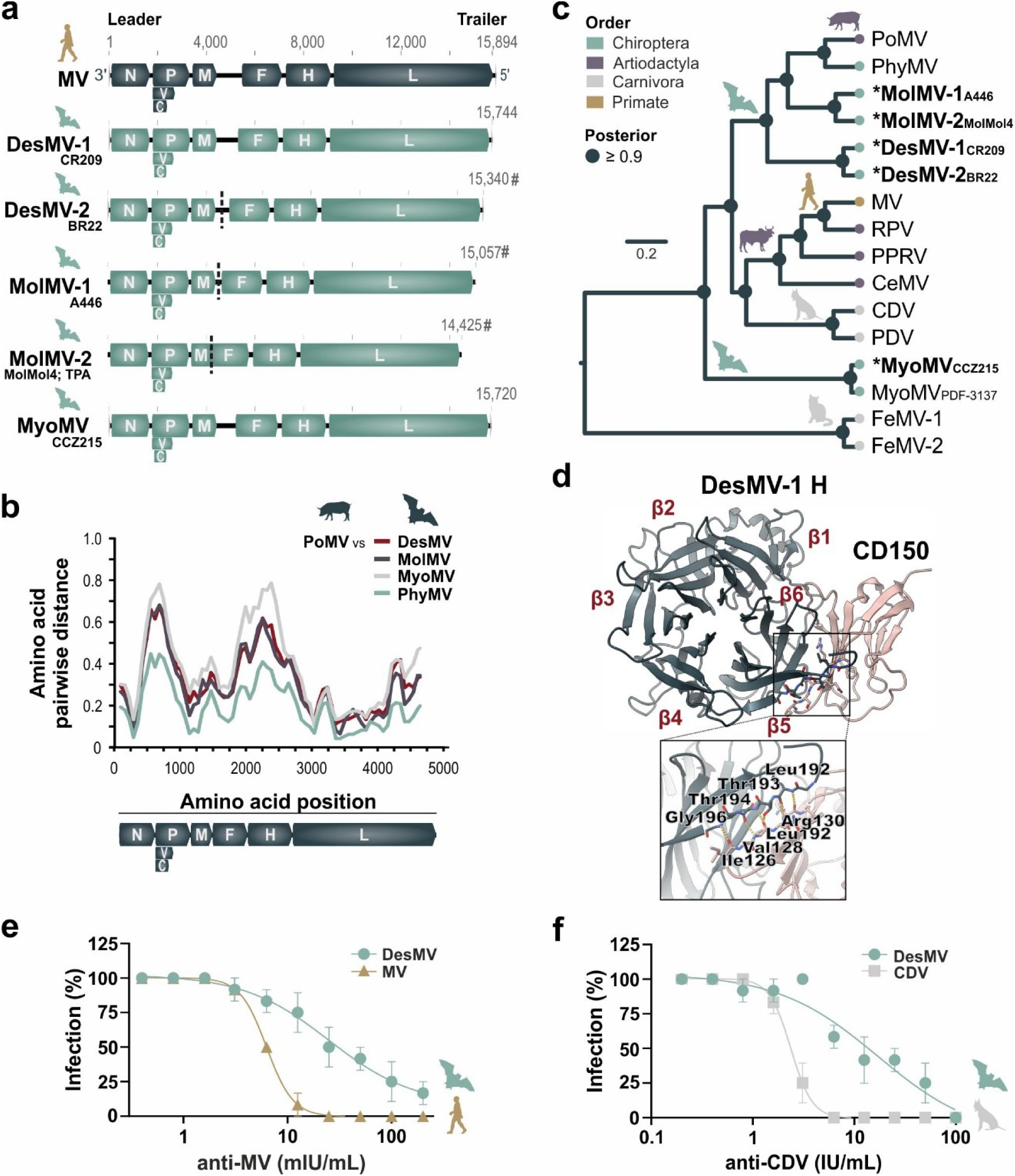
Molecular and *in vitro* characterization of bat-associated morbilliviruses. a) Genome organization of bat morbilliviruses compared to MV. # represents near-complete genome and dashed line represents region with a gap. TPA, third party annotation. b) Comparison of sequence distances of bat-associated morbilliviruses versus PoMV (concatenated translated open reading frames (ORFs)). c) Bayesian phylogeny of morbilliviruses using translated, concatenated ORFs. Symbol **\*** describes sequences from this study. Scale bar indicates genetic distance. See **Supplementary Tables S1-S2** for virus abbreviations and GenBank accession nos. d) Representation of the interaction of DesMV-1 hemagglutinin (H) glycoprotein with cellular receptor CD150. Close-up view shows the interactions between hydrogen-bonding edges of β-sheets of DesMV-1 H glycoprotein and CD150. e) Neutralization curves of DesMV (Vero-vamCD150), MV (Vero-hCD150), and CDV (Vero-76) by anti-MV (upper panel) and anti-CDV sera (bottom panel) measured in international units (IU). Error bars are in standard error of the mean.

Bayesian phylogenetic analysis of translated, concatenated ORFs showed that most bat-associated morbilliviruses characterized on the full-genome level were monophyletic with the exception of MyoMV (**Fig. 2c**). Different tree topologies were reconstructed for the individual ORFs (**Extended Data Fig. 2**), as has been reported for other paramyxoviruses^9^, yet formal analyses (**Supplementary Table S3**) yielded no evidence for recombination involving bat-associated morbilliviruses which is consistent with low rates of recombination in negative-sense single-stranded RNA viruses^24^.

### Conserved antigenicity and entry of bat-associated morbilliviruses

Homology modelling of the receptor binding H glycoprotein of bat-associated morbilliviruses based on MV H glycoprotein co-crystallized with the cellular receptor CD150 (PDB 3LAZ) revealed a typical β-sheet augmentation previously published as an interaction site between MV H glycoprotein and CD150^25^, suggesting conserved usage of CD150 for cellular entry of phylogenetically divergent bat morbilliviruses (**Fig. 2d; Extended Data Fig. 1c-d**), consistent with the isolation of DesMV in Vero-vamCD150 in this study and previous rescue of MyoMV in Vero cells expressing myotis bat CD150^26^.

Cross-neutralization experiments revealed cross-protective immune responses of anti-MV and anti-CDV reference sera against DesMV (**Fig. 2e-f**). As expected, the half maximal inhibitory concentration (IC50) of anti-MV and anti-CDV sera were 4-to 7-fold higher for DesMV compared to MV and CDV, which was comparable to cross-neutralization of a reconstructed bat morbillivirus^26^ and cross-protection of CDV in MV-vaccinated macaques^27^. Infection by a novel morbillivirus may thus be limited by previous morbillivirus-elicited immunity.

### Identification of marmoset-associated morbilliviruses

Relatively high prevalence and diversity of morbilliviruses in Neotropical bats, and existing precedents for virus transmission from bats to non-human primates (NHPs)^28,29^ prompted us to search retrospectively for bat-associated morbilliviruses in Neotropical primates. We investigated liver tissues collected from 1,370 wild NHPs from Brazil representing at least 13 species that died in the wild with suspicions of yellow fever between 2017-2023 (**Table 2**)^30^. All samples included in this study were PCR-negative for yellow fever virus^30^. We detected 13 morbillivirus-positive marmosets by RT-PCR^10^ (*Callithrix jacchus* and *C. penicillata*) from neighboring Brazilian states Bahia and Minas Gerais (**Fig. 3a**, **Supplementary Table S4**) (overall detection rate, 0.95%; 95%CI 0.56-1.62) with viral loads between 10^7^-10^8^ RNA copies/g of liver. Marmoset-associated morbillivirus (MarMV) sequences grouped by location, not by host species, forming two clades differing by about 2% of their genomic sequences (termed Bahia (BA)-1 and -2) and another clade (termed Minas Gerais (MG)) differing by about 5% of its genomic sequence from the Bahia clades (sampling sites had approximately 750 km distance; **Fig. 3b; Extended Data Table 2**). Nearly complete MarMV genomes of about 16 kb lacking only the genome ends were recovered by deep sequencing aided by morbillivirus-targeted in-solution capture. Sequence comparisons showed that MarMV genomes differed from the bat-associated PhyMV^9^ by only 2-5% at the nucleotide level (**Extended Data Table 2**) throughout the genome (**Fig. 3c**). Only the intergenic region between the *M* and *F* genes showed slightly higher divergence (7-12%), which may be accounted for by a six-nucleotide insertion in the PhyMV sequence relative to MarMV genomes. None of the samples for which nearly complete genomes were generated by deep sequencing contained any read that mapped to *cytochrome c* oxidase subunit I of animal species other than *Callithrix* spp. with 99-100% mean identity (**Supplementary html files**) confirming the marmoset host associations of morbillivirus-positive specimens.

**Fig. 3.**
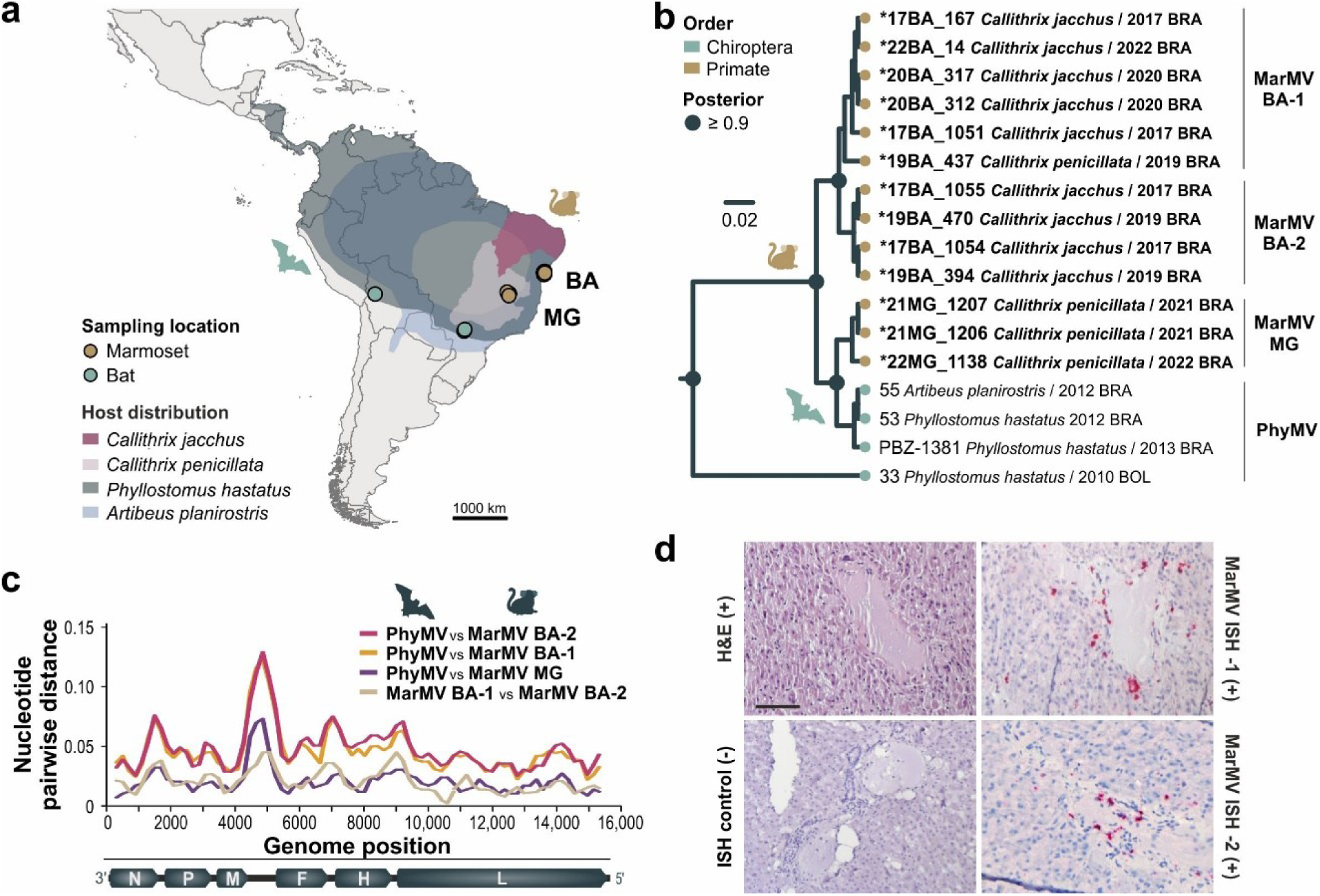
Characterization of marmoset-associated morbilliviruses. a) Geographical distribution of marmoset and bat hosts of related morbilliviruses and sampling locations. BA, Bahia; MG, Minas Gerais. b) Bayesian phylogeny of MarMV and related morbilliviruses using partial *L* gene (361 nucleotides). Symbol **\*** describes sequences from this study. The outgroup used was PoMV. See **Supplementary Tables S1** and **S4** for virus abbreviations and GenBank accession nos. c) Comparison of sequence distances of MarMV representatives of each phylogenetic clade versus PhyMV. d) *In situ* hybridization of MarMV PCR-positive marmoset 17BA_167. Strong multifocal MarMV RNA signal (red staining) is shown throughout the periportal space with endothelial portal vein distribution, and intracytoplasmic signal in the epithelial bile duct and hepatocytes in left panels. No significant histopathological findings were observed in hematoxylin and eosin (H&E) section. No staining was shown in the MarMV-negative animal (ISH control). (+), MarMV-positive animal, (-), MarMV-negative animal. Scale 100µm.

**Table 2.** Non-human primate samples.

| Species | States | Total | Pos | Years (n) |
| --- | --- | --- | --- | --- |
| *Alouatta caraya* | Minas Gerais | 3 |  | 2021 (3) |
|  | Sao Paulo | 1 |  | 2021 |
| *Alouatta guariba clamitans* | Minas Gerais | 2 |  | 2017 (1), 2022 (1) |
| *Alouatta* sp. | Espírito Santo | 2 |  | 2020 (1), 2022 (1) |
|  | Rio de Janeiro | 7 |  | 2017 (2), 2019 (2), 2020 (2), 2021 (1) |
| *Ateles marginatus* | Sao Paulo | 2 |  | 2021 (2) |
| *Ateles paniscus* | Sao Paulo | 1 |  | 2021 |
| *Callithrix aurita* | Sao Paulo | 1 |  | 2021 |
| *Callithrix geoffroyi* | Espírito Santo | 8 |  | 2017 (1), 2018 (1), 2019 (5), 2021 (1) |
|  | Minas Gerais | 7 |  | 2021 (6), 2022 (1) |
| *Callithrix jacchus* | Bahia | 42 | 8 | 2017 (27), 2019 (11), 2020 (2), 2022 (2) |
|  | Minas Gerais | 14 |  | 2021 (10), 2022 (4) |
|  | Rio Grande do Norte | 32 |  | 2018 (3), 2019 (2), 2020 (12), 2021 (9), 2022 (6) |
|  | Sao Paulo | 11 |  | 2020 (7), 2021 (4) |
| *Callithrix penicillata* | Bahia | 38 | 1 | 2017 (34), 2019 (4) |
|  | Minas Gerais | 122 | 3 | 2017 (6), 2021 (43), 2022 (70), 2023 (3) |
|  | Sao Paulo | 25 |  | 2020 (21), 2021 (4) |
| *Callithrix* sp. | Bahia | 132 | 1 | 2017 (48), 2018 (18), 2019 (3), 2020 (11), 2021 (30), 2022 (22) |
|  | Espírito Santo | 34 |  | 2017 (13), 2018 (19), 2020 (2) |
|  | Minas Gerais | 9 |  | 2017 (2), 2021 (3), 2022 (4) |
|  | Rio de Janeiro | 618 |  | 2017 (79), 2018 (270), 2019 (152), 2020 (50), 2021 (47), 2022 (20) |
|  | Rio Grande do Norte | 10 |  | 2020 (1), 2021 (2), 2022 (7) |
|  | Sao Paulo | 9 |  | 2020 (8), 2021 (1) |
| *Callithrix* sp.* | Bahia | 168 |  | 2017 (17), 2018 (43), 2019 (72), 2020 (32), 2021 (4) |
|  | Rio Grande do Norte | 42 |  | 2018 (3), 2019 (18), 2020 (7), 2021 (3), 2022 (11) |
| *Cebus apella* | Rio de Janeiro | 1 |  | 2017 |
| *Cebus* sp. | Minas Gerais | 6 |  | 2017 (2), 2021 (1), 2022 (3) |
|  | Rio de Janeiro | 14 |  | 2017 (2), 2018 (1), 2019 (2), 2020 (3), 2021 (5), 2022 (1) |
|  | Rio Grande do Norte | 3 |  | 2022 (3) |
| *Leontopithecus* sp. | Bahia | 1 |  | 2017 |
| *Pithecia* sp. | Rio de Janeiro | 1 |  | 2021 |
| *Plecturocebus cupreus* | Espírito Santo | 1 |  | 2021 |
|  | Minas Gerais | 1 |  | 2021 |
|  | Sao Paulo | 1 |  | 2020 |
| *Sapajus nigritus* | Sao Paulo | 1 |  | 2020 |
| **Total** |  | **1,370** | **13** |  |

Pos, Morbillivirus positive animals; \* Not morphologically identified, based on most likely species in the region

Attempts to isolate MarMV from 12 available RT-PCR-positive liver samples were neither successful in the tamarin lymphoblastoid cell line B95a traditionally used for morbillivirus isolation (tamarins are genetically related to marmosets), nor in cells expressing human CD150, potentially due to virus degradation during sampling and storage. The analysed livers did not show significant histopathological findings (**Fig. 3d, Extended Data Fig. 3**), suggesting that the MarMV-positive animals did not die of hepatic disease, which is also uncommon in human MV infections^31^. No other marmoset tissues were available to comparatively assess organ tropism, but RNA *in situ* hybridization (ISH) revealed strong positive MarMV *N* gene signal in bile duct epithelial cells and vascular endothelial cells within portal tracts, and in hepatocytes in three MarMV-positive liver tissue samples (**Fig. 3d, Extended Data Fig. 3**), similar to other morbillivirus infections in the liver^32^ and compatible with systemic spread of MarMV.

### Morbillivirus origins in Laurasiatherian hosts

To investigate morbillivirus genealogy, we included all available divergent morbillivirus sequences, including the partially sequenced shrew-associated^12^ and genetically divergent bat-associated morbilliviruses. Most morbilliviral hosts belong to the Laurasiatheria^33^. Bayesian ancestral state reconstruction (ASR; BayesTraits) confirmed the origins of morbilliviruses in Laurasiatherian hosts (log_10_ Bayes factor 6.5, **Extended Data Table 3**). Distance-based coevolutionary analyses were performed in 1,000 trees randomly selected from the posterior distribution of the partial *L* gene-based tree **(Extended Data Fig. 4a)**. An overall association (*p*=0.02±0.01; ParaFit; **Supplementary Table S5**) between morbilliviruses and their hosts was found implying some support for host-virus co-speciation. RNA virus evolution outpaces that of mammalian hosts by orders of magnitude and genetically related strains, which may represent diversification within one viral species clustering apically, like DesMV and FeMV, can confound distance-based coevolutionary analyses. Therefore, we repeated the coevolutionary reconstructions using only one virus per host species (**Extended Data Fig. 4b**). No significant signal for coevolution was detected (*p*=0.10±0.02; ParaFit; **Supplementary Table S5**), consistent with a predominant role of host shifting in morbillivirus macroevolution. We used the three most likely topologies from DensiTree to reconstruct event based phylogenies of historical host shifts (**Fig. 4a** and **Extended Data Fig. 4c**). Host shifts from bats and artiodactyls to other hosts were predominant in the two most likely topologies. Host shifts from carnivores to other hosts were predominant in the third most likely topology. Cophylogenetic analyses in all three most likely topologies consistently reconstructed an origin of PoMV, detected in pigs, from a bat-associated ancestor, an origin of MV from a cattle-associated ancestor^34^, and host switching among bat host species. Parsimony-based ASR showed relatively high average numbers (>0.5) of host shifts from bats to artiodactyls and carnivores, from carnivores to bats and shrews, and from artiodactyls to primates (**Fig. 4b**). Our results imply that bats, carnivores, and artiodactyls are main sources of cross-order host shifts in morbilliviruses. However, the probability of a bat origin was consistently highest in all major nodes [posterior probability (PP) 0.30-0.95 in nodes A-D, **Fig. 4c**]. Removing morbilliviral genomic sequences generated in this study led to lower statistical support of bat origins (0.24 – 0.60 in nodes A-D, **Extended Data Fig. 5**). Although shrew morbilliviruses clustered in basal relationship to most other morbilliviruses, they formed a monophyletic clade (**Fig. 4c**) and showed low average numbers of reconstructed host shifts (<0.2) (**Fig. 4b**), suggesting that shrews are ancient hosts of morbilliviruses that may not be involved in cross-order host shifts to the same extent as other host orders. Thus, the evolutionary origin of morbilliviruses was strongly associated to Laurasiatherian hosts in general and bats in particular.

**Fig. 4.**
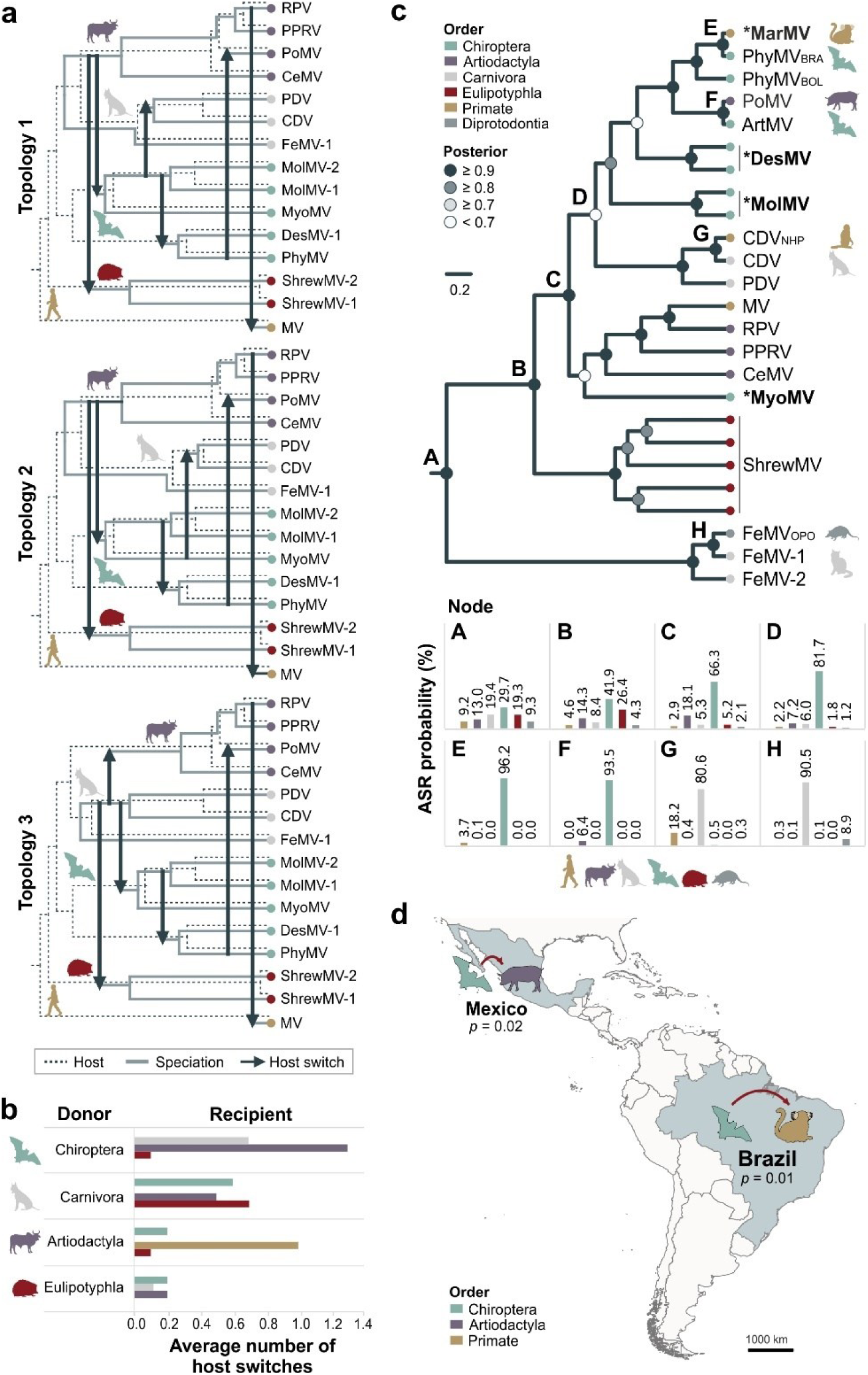
Macroevolution of morbilliviruses. a) Event-based coevolutionary analyses of the three most likely topologies: 1- most likely, 2- second most likely, 3- third most likely. b) Average number of morbillivirus host switches from parsimony-based ancestral state reconstructions (ASR). c) *Upper* insert, Bayesian phylogeny of representative morbillivirus species and sequences from suspected cross-order spillover events. Symbol **\*** describes sequences from this study and scale bar indicates genetic distance. BOL, Bolivia; BRA, Brazil; NHP, non-human primate; OPO, opossum. *Lower* insert, posterior probabilities for Bayesian ancestral state reconstructions (ASR) for each evaluated node (A-H) are shown as a graph. See **Supplementary Tables S1-S2 and S4** for virus abbreviations and GenBank accession nos. d) Graphical representation of the geographic association of bat-, marmoset-, and pig-associated morbilliviruses; *p* values indicate the significance of the association.

### Evidence for cross-order spillover events from bat-associated morbilliviruses

Cross-order transmissions have been reported for CDV, including to macaques (order Primates) and collared peccaries (New World pigs, order Artiodactyla)^35^. Other reported cross-order morbillivirus transmissions include the detection of FeMV in opposums^36^ (order Diprotodontia), and CeMV in seals^37^ (order Carnivora). To test if bat-associated morbilliviruses are main sources of cross-order host shifts, we analysed morbilliviruses identified in a different host order than the putative reservoir host, including the MarMV described above (**Figure 4c**, *upper insert*). Bayesian ASR at those nodes (**Figure 4c**, *lower insert*) suggested a bat origin for MarMV (PP>0.96, node E) and PoMV (PP>0.93, node F), as well as a carnivore origin for macaque-CDV (PP>0.81, node G) and opossum-associated FeMV (PP>0.91, node H). Testing in a Bayesian framework corroborated geographic association (*p*<0.05; BaTS) for bat- and marmoset-associated morbilliviruses from Brazil as well as the bat- and pig-associated morbilliviruses from Mexico (**Fig. 4d, Extended Data Table 4**). Notably, bat-associated morbillivirus sequences were neither identified in the genomic data from 82,178 samples of NHPs nor in 102,427 samples of artiodactyls in the NCBI SRA repository. The lack of morbillivirus sequence data in those animals may suggest repeated bat-to-NHP and a bat-to-pig transmission as compared to maintenance of bat-associated morbilliviruses in those animals after an initial host shift.

## Discussion

Our macroevolutionary data project the origins of major morbilliviruses to non-recent host shifts, predominantly involving bat hosts as source. Bats are also important reservoirs of other human-pathogenic paramyxoviruses, including henipa- and rubulaviruses^10^. A prominent exception to the relevance of bats as sources of morbillivirus host shifts according to our and published data^34^ is the projected origin of human MV in an ancestor carried by cattle. However, this hypothesis is largely based on the close genetic relationship of MV and RPV. According to the available genomic data and our macro-evolutionary reconstructions, neither cattle nor other ungulates, as opposed to bats, are reservoirs of genetically divergent morbilliviruses that have caused cross-order host switches. Whether MV indeed originated from an ancestor carried by cattle or from an alternative reservoir, thus requires urgent investigation.

The Neotropics are a hotspot for bat diversity and predicted zoonotic richness from bats^38^. All bat morbilliviruses so far were found in the Neotropics, supporting the anecdotal evidence on the New World origin of CDV, first reported in Peruvian dogs in 1746^22^. A Neotropical origin of major animal morbilliviruses was also consistent with the likely bat origin of PoMV^8^. Contrastingly, MV and RPV likely emerged more recently according to apical clustering and relatively short branch in the morbillivirus phylogeny^34^, and are suggested to have an Old World origin according to the earliest descriptions of measles in Asia around 900 CE and of the introduction of rinderpest to Europe during the Hun invasion around 400 CE^2,22^. This interpretation is consistent with lethal MV outbreaks in morbillivirus-naïve indigenous populations during the European colonization of the Americas^22^. However, intense surveillance of Old World bats has not yielded positive results for morbilliviruses^9,10,39^, including the mining of sequence data in this study. Whether morbillivirus evolutionary origins date back to Neotropical bat dispersal histories (40 -13 million years ago)^40–42^ and extinction of morbillivirus in Old World bats, or to macroevolutionary phenomena, such as preferential host switching in sympatric animals or via intermediate hosts, or whether those estimates are affected by sample bias remains to be investigated. It is possible that zoonotic events from Neotropical bats and other Neotropical animal reservoirs of human pathogenic viruses are underappreciated, requiring increased surveillance and funding for research and surveillance. Targeted field work potentially elucidating the timing and hosts of morbilliviral macro-evolution should include Neotropical shrews and Old World bat species genetically related to Neotropical bats hosting morbilliviruses.

Potential spillover events from bat-associated morbilliviruses into livestock may be facilitated by intense rearing of cattle and pigs in Latin America, including both Brazil and Mexico^43,44^. Land use changes such as agricultural encroachment in natural habitats may further facilitate spillover events from bats to livestock^45^, reminiscent of outbreaks of the bat-borne Nipah virus and a bat-associated coronavirus in pigs in Asia^45,46^. Predator-prey interactions may be another potential cross-order host shift driver in morbilliviruses, as suggested for carnivore-associated morbilliviruses^22^, including feeding of vampire bats on diverse mammals, including animals reared as livestock, pets and humans^47^. Spillovers of bat-associated viruses to humans and NHPs have been reported for other viruses in Latin America, such as RABV^28,48^. Marmoset feeding of bat-contaminated fruits may be a plausible transmission route, as proposed for the transmission of Marburg virus to apes in Africa^29^. Transmission to pigs may occur similarly from feeding of bat-contaminated fruits, as reported for Nipah virus, which can also transmit directly to humans via consumption of bat-contaminated date palm sap^49^. In parts of Latin America, bat consumption for traditional medicine may also be a potential morbillivirus transmission route^50^.

Limitations to our study include sampling in only two Neotropical countries. However, this was accounted for by performing data mining of publicly available genomic sequences. Our macroevolutionary reconstructions were limited by phylogenetic uncertainty due to usage of partial *L* gene sequences of genetically divergent bat morbilliviruses. However, this was accounted for by using different phylogenetic methods and tree topologies. Additionally, lack of recombination in morbilliviruses enhances generalizability of *L* gene-based phylogenies compared to other viral taxa.

Evidence for bat-to-pig spillover leading to PoMV emergence suggests that bat morbilliviruses may be immediate threats for morbillivirus-naïve livestock, such as cattle and pigs. Hypothetically, partial cross-protective immunity against heterologous morbilliviruses may enable cost- and time-effective mitigation strategies relying on vaccines licensed for use in livestock against rinderpest and peste des petits ruminants (PPR). However, due to the cessation of rinderpest vaccination in the early 2000s^51^, rinderpest vaccine stocks are limited today to only four facilities in the world, located in Asia, Africa and Europe^52^. Additionally, whether rinderpest and PPR vaccines confer effective protection against heterologous morbilliviruses^53^ or whether PPR vaccines are suitable for use in pigs is not well known^54^. New vaccines for use in those animals reared as livestock for human consumption in case of further outbreaks of reservoir-borne morbilliviruses will need time and may be technically challenging^55^. However, development and licensing of specific vaccines against emerging viruses within 100 days is envisaged in programs such as CEPI 2.0 (https://cepi.net/) and may have to consider pathogens threatening both humans and other animals. Finally, our finding of bat-associated morbillivirus host shifts into NHPs suggest that once MV is eliminated, termination of mandatory or recommended MV vaccination may lead to outbreaks caused by reservoir-bound morbilliviruses in humans, reminiscent of mpox emergence after smallpox elimination^56^. Thus, our data advocate for intensified surveillance, experimental risk assessments, and vaccine development for reservoir-bound morbilliviruses.

## Online Methods

### Animal sampling

Bat sampling was performed using mist nets at roost or foraging sites at different locations in Costa Rica and Brazil (**Supplementary Table S2**). After capturing, animals were identified, anesthetized, euthanized and dissected immediately by a trained field biologist. The euthanasia of the bats followed the AVMA 2020 Guidelines for the Euthanasia of Animals. Samples were stored in separate tubes with and without RNAlater for each organ at -20°C 2 to 3 days before being transferred to -80°C. We also included bats that were found dead in the wild and were sent to Rabies surveillance laboratories in Brazil. In Brazil, authorizations were granted by ethics committee of the Universidade Estadual Paulista (UNESP) (009/2012), Instituto Ambiental do Paraná (235/10), the ethics committee of the Institute of Biomedical Science from the University of São Paulo (56–18–03/2014), Ministry of the Environment of Brazil (51231-1), and the Instituto Brasileiro do Meio Ambiente e dos Recursos Naturais (12751-2, 12751-3, 21748-1, 27346-2). In Costa Rica, the capture of vampire bats is part of the National Surveillance Bovine Rabies Program by the Animal Health Service (SENASA) following the Executive Decree 24405-MAG of June 8, 1995, of the Ministry of Agriculture of Costa Rica in which the control of vampire bats is declared compulsory. All procedures were carried out according to the “Ley de Bienestar de los Animales” (Ley No 7451 1994) and the International Convention for the Protection of Animals endorsed by Costa Rican Veterinary Law on the CR-NAHS (Ley No 8495 2006). Some of the samples used in this study have been previously described (**Table 1**).

All NHP samples analysed in this study were submitted for flavivirus surveillance programs to the Reference Laboratory for Arboviruses and Hemorrhagic Viruses at the Oswaldo Cruz Institute in Brazil between 2017-2023 (**Supplementary Table S4**). The study was approved by the institutional research ethics board under protocol IOC 274/05. All samples previously tested negative for yellow fever virus by a real-time RT-PCR^30^. CITES export permits for the NHPs were granted by The Brazilian Institute of the Environment and Renewable Natural Resources (IBAMA): 23BR046601/DF. CITES import permits were granted by the German Federal Agency for Nature Conservation: DE-E-02733/23-DE-E-02736/23, and sample import permits were granted by the Senate Department for Justice and Consumer Protection: EG I – 2023/62.

Animal species of positive samples were confirmed by a nested PCR targeting the taxonomic marker *cytochrome c* oxidase subunit I (COI) gene as previously described^57^. In addition, to confirm that no other animal species was present in the analysed samples, all host reads were mapped to a non-redundant database of COI sequences from GenBank using Bowtie2^58^. Next, a sequence similarity search was performed for the mapped reads using the blastn algorithm. Bat species distribution were retrieved from IUCN Red List of Threatened Species (www.iucnredlist.org) and mapped using R v3.4.1 through the Rstudio environment v4.0.3.

### Sample preparation and virus detection

Frozen samples were homogenized in the Tissue Lyser (Qiagen). RNA was extracted from homogenized material using MagNA Pure 96 Viral NA Large Volume Kit (Roche Molecular Systems, USA). Samples were then screened for the presence of paramyxovirus RNA with a broadly reactive semi-nested reverse transcription (RT)-PCR targeting the polymerase gene of the family *Paramyxoviridae* (PAR) and the genera *Respirovirus*, *Morbillivirus*, and *Henipavirus* (RMH) as previously described^59^ using a SuperScript III One-Step kit (Thermo Fisher Scientific, Massachusetts, USA). Positive bands were sent for Sanger sequence (Microsynth Seqlab, Germany). After obtaining the genetic sequences, quantitative RT-(q)PCR were designed targeting the novel sequences using newly designed primer sets (IDT Technology; **Supplementary Table S6**).

### Sequencing and genome retrieval

The genome of sample BR22 from a vampire bat from Brazil^10^ was generated by multiple RT-PCRs targeting conserved regions of morbilliviruses CDV, PDV, MeV, RPV, PPRV and CeMV (**Supplementary Table S7**). For all other samples, a high throughput sequencing approach was performed. RNA libraries were prepared for the positive samples according to KAPA HyperPrep manufacturer protocol (Roche) for sequencing on a NovaSeq 6000 system (250 cycles paired-end). Low quality reads of <Q30 were trimmed with Trimmomatic V.0.4. After quality trimming, remaining reads were aligned to a morbillivirus protein reference database (Ensemble) using DIAMOND (https://github.com/bbuchfink/diamond). Aligned reads were extracted and *de novo* assembled using Geneious assembler within Geneious R.11.1.5 (https://www.geneious.com). A reference-based assembly using all high-quality reads was then performed with the newly generated morbillivirus contigs using Bowtie2. Several iterations were performed until no more new reads were assembled. A morbillivirus-targeted capture library was also used for bat-associated morbillivirus-positive marmoset samples following the manufacturers’ protocol (BioCat, Germany). Morbillivirus and morbillivirus-related sequences included in the capture library are described in **Supplementary Table S8**. The generated capture-based libraries were sequenced on a MiniSeq system (150 cycles paired-end). Quality trimming was performed as indicated above. Reads were mapped to the associated bat morbillivirus using Bowtie2 to generate complete genomes. Gaps in the coding region were filled by PCR and Sanger sequencing (Microsynth Seqlab). Robustness of all assembled genomes from Illumina HTS, including the MolMV recovered from third party annotation, was confirmed by performing a reference assembly with a minimum mapping quality of 30 and allowing only ≤1 % mismatches per read to the consensus genome using Geneious algorithm. Regions of low coverage (<2) were confirmed by Sanger sequencing. Median coverage for genomes ranged from 10.5 – 12,372.1 (**Extended Data Fig. 6**). To obtain genome ends, we performed a rapid amplification of cDNA ends (RACE) according to the 5’/3’ RACE kit protocol (Roche) with modifications. Briefly, RNA was converted to cDNA using primers targeting both 5’ and 3’ followed by column purification using Monarch PCR & DNA cleanup kit (New England Biolabs, Massachusetts, USA). A poly(A) tail was then added using dATP and terminal transferase. Thereafter, in separate reactions for each genome termini, we performed a first round of PCR using a second set of primers targeting either 5’ or 3’ and an oligo dT-anchor primer from the kit. After amplification, product purification by column was performed and a nested PCR was set up using a third set of primers targeting either 5’ or 3’ and a PCR anchor primer from the kit. Products were sent for Sanger sequencing (Microsynth Seqlab). Only the 5’ RACE was used to determine both genome termini as both genome and anti-genome are present in organs in which virus replication takes place.

### Data mining

We searched for unidentified divergent morbilliviruses by using the web tools Serratus explorer and palmID: Viral-RdRP Analysis (https://serratus.io/toolkit). Both tools rely on data mining of the SRA sequencing libraries. Once a morbillivirus sequence was identified, the SRA sequence was downloaded and *de novo* assembly was performed. Contigs were BLASTXed against NCBI non-redundant database. Morbillivirus contigs were used as the reference to do reference-based assemblies using Bowtie2 with various iterations until no more new reads were mapped. The mammal diversity database v1.10 (10.5281/zenodo.7394529) of the American Society of Mammalogists was used to gather taxonomic information and the geographic distribution of the mammals in the SRA dataset used in the Serratus platform.

### Data set

Morbilliviruses were chosen based on ICTV morbillivirus species. Only one representative sequence of a morbillivirus species was included. Virus species was defined according to ICTV species demarcation criteria: branch lengths of >0.3 in the L protein. We excluded diverse rodent paramyxoviruses that occur in phylogenetic sister relationship to morbilliviruses from our analyses as these sequences had a different genome organization and lacked conserved key amino acids in H glycoprotein for CD150 binding^60^. These rodent-borne viruses may thus have had a different evolutionary pathway to classical morbilliviruses. Sequences used in this study were downloaded from GenBank via Geneious. The dataset consists of publicly available genome sequences that were accessed until 8^th^ of July 2022. Divergent morbilliviral sequences in bat hosts published thereafter were also included. Genome sequences used for evolutionary analyses are provided in **Supplementary Table S1**.

### Protein modelling

All hemagglutinin (H) models of the bat morbilliviruses were generated by homology modelling using Swiss model engine (www.swissmodel.expasy.org/) based on the MeV H and tamarin CD150 structures from PDB 3ALZ^25^ as template using default settings. Models were superimposed using least-square fit onto the hemagglutinin of 3ALZ chain E in ChimeraX 1.0 (https://www.rbvi.ucsf.edu/chimerax/). Root mean square deviations (RMSD) of the backbone atoms were calculated based on a C-alpha atom alignment using ChimeraX 1.0.

### Cells, plasmid, and viruses

All cells were cultured in DMEM media supplemented with 10% FBS, and 1% penicillin/streptomycin (PS). Cells used were Vero 76 (CCLV-RIE 228), Vero-hCD150 (Vero-hSLAM) that express human CD150, and Vero-vamCD150 that express vampire bat CD150. Vero-vamCD150 cells were produced by transfecting a plasmid containing vampire bat CD150 (pCAGGS-IgK-HA-vamCD150) by FuGENE® HD and multicycle puromycin selection. The plasmid was generated as previously published^61^. In brief, we subcloned a synthetic fragment containing the immunoglobulin IgK leader sequence, followed by HA epitope with a linker sequence to the vampire bat CD150 ORF (GenBank accession no. XM_024571326.1), for which we removed the signal sequence (amino acids 1-28) predicted by signal P 4.1 software (www.cbs.dtu.dk/services/SignalP/). Viruses used include CDV (Onderstepoort), MV (D8), and DesMV (CR209). Vero 76 cells and CDV were kindly provided by Sven Reiche from Friedrich-Loeffler-Institut, Germany, MV was kindly provided by Annette Mankertz from Robert Koch Institute, Germany, Vero-hCD150 were kindly provided by Yusuke Yanagi from Kyushu University, Japan, and the B95a cells were kindly provided from Rik de Swart from Erasmus MC Rotterdam, Netherlands. Cell line aliquots were sent regularly for qPCR-based mycoplasma testing during experiments (Eurofins Genomics, Germany). All cell lines tested negative during the experiments.

### Virus isolation and kinetics

DesMV-positive sample homogenates were isolated in Vero-vamCD150 cells. MarMV-positive sample homogenates were isolated in B95a and Vero-hCD150 cells. Successful isolation was evaluated by monitoring the viral load by RT-qPCR. The virus titer for DesMV isolate was determined by TCID_50_ assay in Vero-vamCD150 cells. The wells were scored as positive when cytopathic effect (CPE) or syncytia formation was observed. DesMV growth kinetics were evaluated in Vero-vamCD150, Vero-hCD150 and Vero 76 cells. Cells were seeded into 12-well plates and inoculated with DesMV at a multiplicity of infection (MOI) of 0.01. Cells were washed twice with PBS and 1.2 mL of DMEM media supplemented with 1% FBS, and 1% PS was added. Supernatants from triplicate wells of each cell line were taken every day for five days and stored at -80°C. Viral load was quantified by TCID_50_ assay and RT-qPCR.

### Microneutralization assay

Serial dilutions of the international standards anti-measles serum (97/648, NIBSC) and anti-canine distemper serum (CDS, NIBSC) were incubated with 100 TCID_50_ of DesMV, MV, and CDV for 1 h at 37°C in quadruplicate wells. Thereafter, mixes of virus and sera were added to Vero-vamCD150, Vero-hCD150 and Vero 76 cells respectively and incubated at 37°C for 5-7 days. Cells were fixed in formalin and stained with crystal violet. Three independent experiments were performed. Virus infection was determined by the presence of CPE, which indicated that no neutralizing activity was generated by antibodies. Wells with CPE were assigned as positive, whereas wells with no CPE were assigned as negative. The percentage of infection was calculated from the number of positive wells at each concentration of antibody. The IC50 values were calculated using the model Nonlinear fit [Inhibitor] vs. response (four parameters) and neutralization curves were generated in GraphPad Prism v9.5.1 software.

Vampire bat sera obtained from field work in Costa Rica in 2016 and 2020 were tested for the detection of antibodies against DesMV. In brief, two-fold dilutions of serum starting at 1:25 were pre-incubated with 100 TCID_50_ of DesMV for 1 h at 37°C. Thereafter, mixes of virus and sera were added to Vero-vamCD150 and incubated at 37°C for 7 days. Cells were fixed in formalin and stained with crystal violet. Neutralizations were conducted in duplicates. Neutralization titers were determined by assigning the dilution in which CPE appeared.

### *In situ* hybridization

Frozen liver samples from MarMV PCR-positive marmoset samples 17BA_1051, 17BA_1054, and 17BA_1055 were thawed and fixated in 4% neutral-buffered formalin. Samples were then sent to iPATH.Berlin at Charité - Universitätsmedizin Berlin, for paraffin embedding, haematoxylin and eosin staining and *in situ* hybridization (ISH) evaluation. A probe targeting specific PhyMV N ORF was designed by Advanced Cell Diagnostics (Hayward, California, USA). ISH was performed using RNAscope 2.5 HD reagent kit-RED assay (Advanced Cell Diagnostics, Inc.) following manufacturer instructions for FFPE samples. The section was counterstained with haematoxylin and mounted with Ecomount. Negative controls consisted of marmoset (*Callithrix jacchus*) liver samples from Costa Rica negative for PhyMV. Of note, the MarMV-positive liver samples analysed may have been decomposed before sampling and may have gone through several cycles of freeze-thaw as all marmosets were found dead in the wild before arriving to the Reference Laboratory for Arboviruses and Hemorrhagic Viruses at the Oswaldo Cruz Institute in Brazil, therefore impacting the structure and architecture of the liver.

### Evolutionary analyses

Sequence alignments were performed with MAFFT v7.450 plugin with an iterative refinement algorithm G-INS-I implemented within Geneious. Mean pairwise sequence distances were calculated using MEGA-X. Sequence distances were calculated and visualized with the Simple Sequence Editor (SSE) v1.4 platform using a sliding window of 600 and step size of 200 amino acids. Neighbor-joining trees were generated based on p-distance model using 1000 bootstrap replicates with MEGA-X. The phylogenetic tree of the complete L protein sequence was generated by Maximum Likelihood method using the LG protein substitution model and 1000 bootstrap replicates in MEGA-X. Recombination analyses were performed for concatenated complete ORFs of morbilliviruses with the methods RDP, GENECONV, Chimaera, MaxChi, BootScann, SiScan, and 3Seq using default parameters in package RDP4 v4.101^62^. Recombinant sites that were detected with >3 methods and p<0.05 were considered as recombination event. Sites with ambiguous data and gaps were excluded from the final alignments used for Bayesian phylogenies. Bayesian phylogenies for translated concatenated ORFs and individual ORFS were generated with BEAST package v1.10.4 (https://beast.community/) using LG+F+I+G4 as substitution model, an uncorrelated lognormal relaxed molecular clock model, and as tree prior the Yule speciation process. The analyses were run for 10 million generations with 10% burn-in, sampling every 1000 steps. A run was considered to have reached convergence when the effective sample sizes of all parameters were >200. Convergence was assessed in Tracer v.1.7.1 (https://beast.community/tracer). The analyses were run for 10 million generations with 10% burn-in, sampling every 1000 steps. A run was considered to have reached convergence when the effective sample sizes of all parameters were >200. Convergence was assessed in Tracer v.1.7.1 (https://beast.community/tracer). The selection of the substitution model was based on the best-fit model for complete and partial L for genus *Morbillivirus* generated by ModelFinder implemented in IQ-TREE. The outgroup used was Longquan Berylmys bowersi morbillivirus 1 (GenBank accession no. MZ328284).

Coevolutionary reconstructions were performed using partial *L* gene sequences (528nt) of representative virus species of the genus *Morbillivirus*. Due to the short sequences used to generate the phylogenetic tree, we generated both nucleotide- and amino acid-based trees in using BEAST package v1.10.4. We used as substitution models GTR+F+I+G4 for the nucleotide-based tree, and LG+F+I+G4 for the amino acid-based trees. Other priors were set as described above. Distance-based cophylogeny was tested in 1,000 randomly selected parasite trees using ParaFit (https://github.com/cran/ape/blob/master/R/parafit.R) was run in R v3.4.1 through the Rstudio environment v4.0.3, with the packages APE v4.1 and Vegan v2.6-2, with 10,000 random permutations of virus-host associations included to test for statistical significance. Event-based cophylogenetic analyses were performed using eMPRess v1, a systematic cophylogeny reconciliation tool^63^. The event costs were fixed to 0 for speciation, 1 for sorting, 2 for duplication and 3 for host switch. The three most likely topologies (blue, red, and green) as indicated by DensiTree^64^ were used to account for phylogenetic uncertainties. The host tree used for cophylogenetic analyses was based on cytochrome B sequences (**Extended Data Fig. 4d**) and according to the mammal tree^33^. Geographic association was tested using BaTS v.0.9^65^, in which the discrete character states represented by the country of origin of the sample. Ancestral state reconstructions (ASR) in a Bayesian framework to evaluate the probabilities between Laurasiatherian and Euarchontoglires hosts at the root were generated using the amino acid-based tree with BayesTraits v3.0^66^. ASR using parsimony method were performed in Mesquite v3.81 as described previously^10^. ASR in a Bayesian framework to evaluate the probabilities of each host order at different nodes was performed using partial *L* gene sequences (361nt) of representative virus species of the genus *Morbillivirus* and sequences from suspected cross-order spillover events in BEAST package v1.10.4.

## Ethics and inclusion statement

This study was performed in collaboration with universities and institutes in Brazil, Costa Rica, and Germany. All participants that made substantial contributions involving project specific fieldwork, had project specific funding, generated and analysed data, wrote and substantially revised the work were included as co-authors, those who designed and performed fieldwork were included in the bat morbillivirus consortium, and those who made minor contributions were included in the Acknowledgements. All permits are described in the Animal Sampling section.

## Data availability

Sequences generated in this study were uploaded to GenBank (accession nos. OP628141-OP628170, PP059213-PP059216, PP850113-PP850121). Assembled genomes and mapped reads were uploaded to NCBI SRA (BioProject ID PRJNA1121230, SAMN41740113-SAMN41740123). Source data are provided in this paper.

## Code availability

Script used for distance-based coevolutionary analyses is available on GitHub (https://github.com/drexler-virus-epidemiology/parafittreesamplerR).

## Acknowledgments

We thank Sebastian Brünink, Nadine Olk, and Arne Kühne for technical support. We thank Christian Drosten for critical reading of the manuscript. We thank Anja Kühl from iPATH.Berlin at Charité - Universitätsmedizin Berlin for technical assistance during ISH. We thank Victor Hugo Sancho for field assistance in Costa Rica. We thank Laura Bergner for assistance on the Serratus Explorer platform usage. This study was supported by the Human Frontier Science Program (grant agreement no. RPG0013/2018), the Defense Advanced Research Projects Agency (DARPA) Prophecy Program award (HR0011-13-2-0020), the Global Centres for Health and Pandemic Prevention from the German academic exchange services (DAAD) (Grant agreement: 57592642 - GLACIER), the Brazilian National Council for Scientific and Technological Development – CNPq (grant agreement no. 203084/2019-5), and the São Paulo state Research Foundation (FAPESP) (grant agreement nos. 2013/11.006-0, 2014/15090-8 and 2017/20744-5), and the University of Costa Rica (Pry01-1681-2019). D.G.S. was funded by a Wellcome Trust Senior Research Fellowship (217221/Z/19/Z) and the UK Medical Research Council (MC_UU_00034/3).

## Competing Interest Statement

The authors declare no conflict of interest.

## Bat Morbillivirus Consortium

Antje Kamprad^1^, Murilo Henrique Anzolini Cassiano^1^, Alejandro Alfaro Alarcon^1,11^, Rocío González-Barrientos^12,13^, Bernal Leon^12^, Cristiano de Carvalho^14^, Wagner André Pedro^14^, Luzia Helena Queiroz^14^, Adriana Ruckert da Rosa^15^, Mateus de Souza Ribeiro Mioni^16,17^, Jane Megid^16^, Andreas Stöcker^18^, Aroldo José Borges Carneiro^19^, Carlos Roberto Franke^19^, Edison Luiz Durigon^20,21^

^11^ Department of Pathology, School of Veterinary Medicine, National University, Heredia, Costa Rica

^12^ Biosecurity Laboratory, Servicio Nacional de Salud Animal (SENASA), LANASEVE, 3-3006 Heredia, Costa Rica

^13^ Texas A&M Vet Med Diagnostic Lab, College Station, Texas 77843, USA

^14^ Dept. de Apoio, Produção e Saúde Animal, Faculdade de Medicina Veterinária de Araçatuba, Universidade Estadual Paulista “Júlio de Mesquita Filho” (UNESP), 16050-680 Araçatuba, São Paulo, Brazil

^15^ Divisão de vigilância de zoonoses de São Paulo, Coordenadoria de vigilância em saúde, São Paulo 01221-010, São Paulo, Brazil

^16^ Department of Animal Production and Preventive Veterinary Medicine, Faculty of Veterinary Medicine and Animal Science, São Paulo State University “Júlio de Mesquita Filho” (UNESP), Botucatu 18618-681, São Paulo, Brazil

^17^ Department of Pathology, Reproduction and One Health, School of Agricultural and Veterinarian Sciences, Sao Paulo State University “Júlio de Mesquita Filho” (FCAV/UNESP), Jaboticabal 14884-900, São Paulo, Brazil

^18^ Infectious Diseases Research Laboratory, University Hospital Professor Edgard Santos, Federal University of Bahia, 40110-060 Salvador, Brazil

^19^ School of Veterinary Medicine, Federal University of Bahia, 40170-110 Salvador, Brazil

^20^ Institute of Biomedical Sciences, University of São Paulo, 05508-900 São Paulo, Brazil

^21^ Institut Pasteur de São Paulo, São Paulo 05508-020, São Paulo, Brazil

---

**Extended Data Fig. 1.**
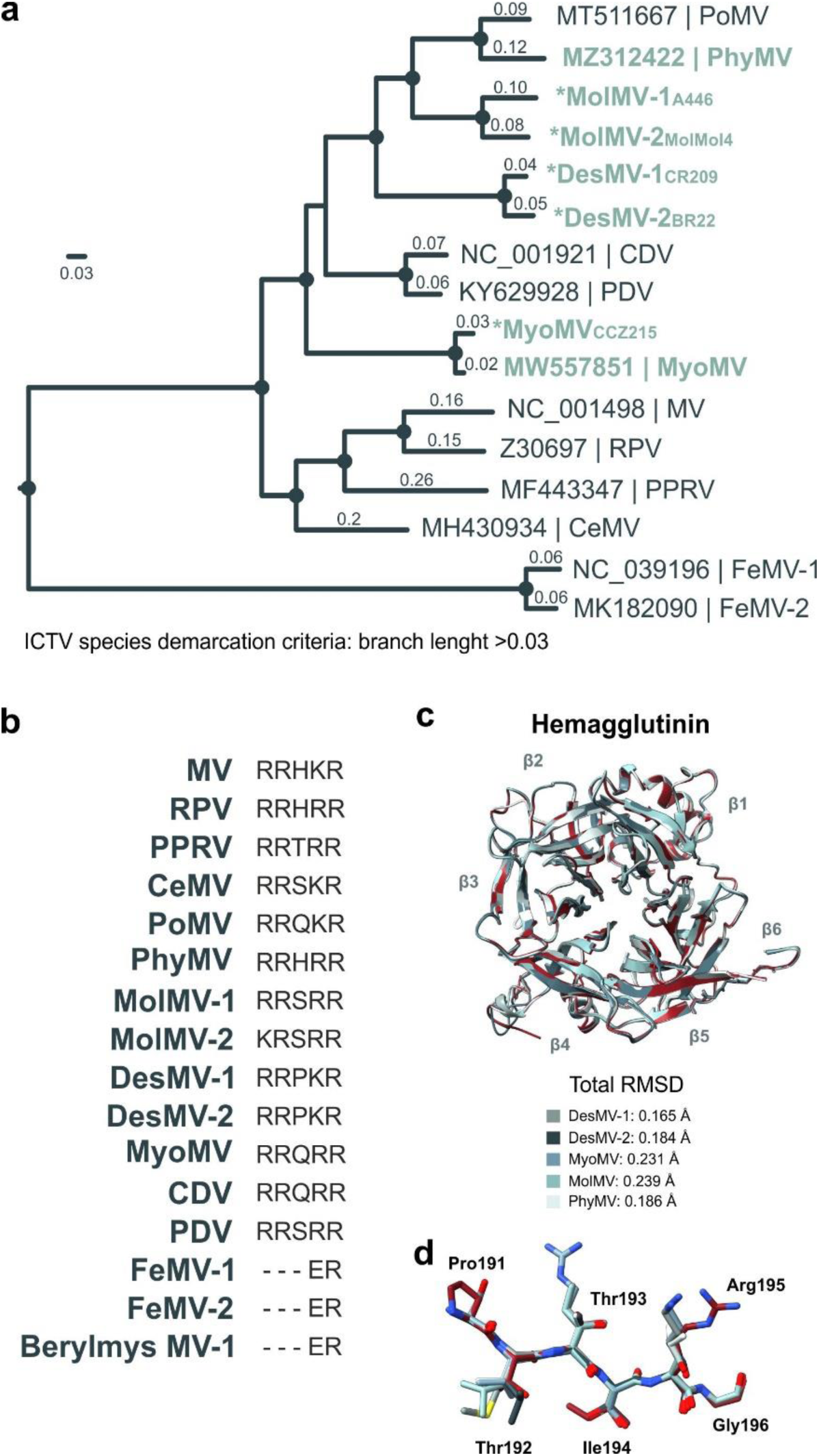
Characterization of morbilliviruses. a) Species demarcation of morbilliviruses based on phylogenetic tree of complete morbillivirus polymerase protein-coding sequences. Tree was inferred by Maximum Likelihood method using LG as protein substitution model with 1000 bootstrap replicates. Circles at nodes indicate bootstrap values ≥80%. Branch lengths (genetic distance) are indicated at each branch. Scale bar indicates genetic distance. b) Polybasic Furin cleavage signal in the F glycoprotein. c) Hemagglutinin structure models of bat morbilliviruses based on MV hemagglutinin (H) complex with tamarin CD150 (PBD 3ALZ). Representation of superimposed hemagglutinin glycoprotein structures of bat morbilliviruses and MV H (red). d) Representation of superimposed beta-sheet augmentation in interaction site 3 of bat morbilliviruses and MV (red). Polypeptide backbones are based on the Pro191–Gly196 strand of MV-H (β6 sheet). RMSD, root mean square deviation; Å, Angstrom. See **Supplementary Tables S1** and **S2** for virus abbreviations and GenBank accession nos.

**Extended Data Fig. 2.**
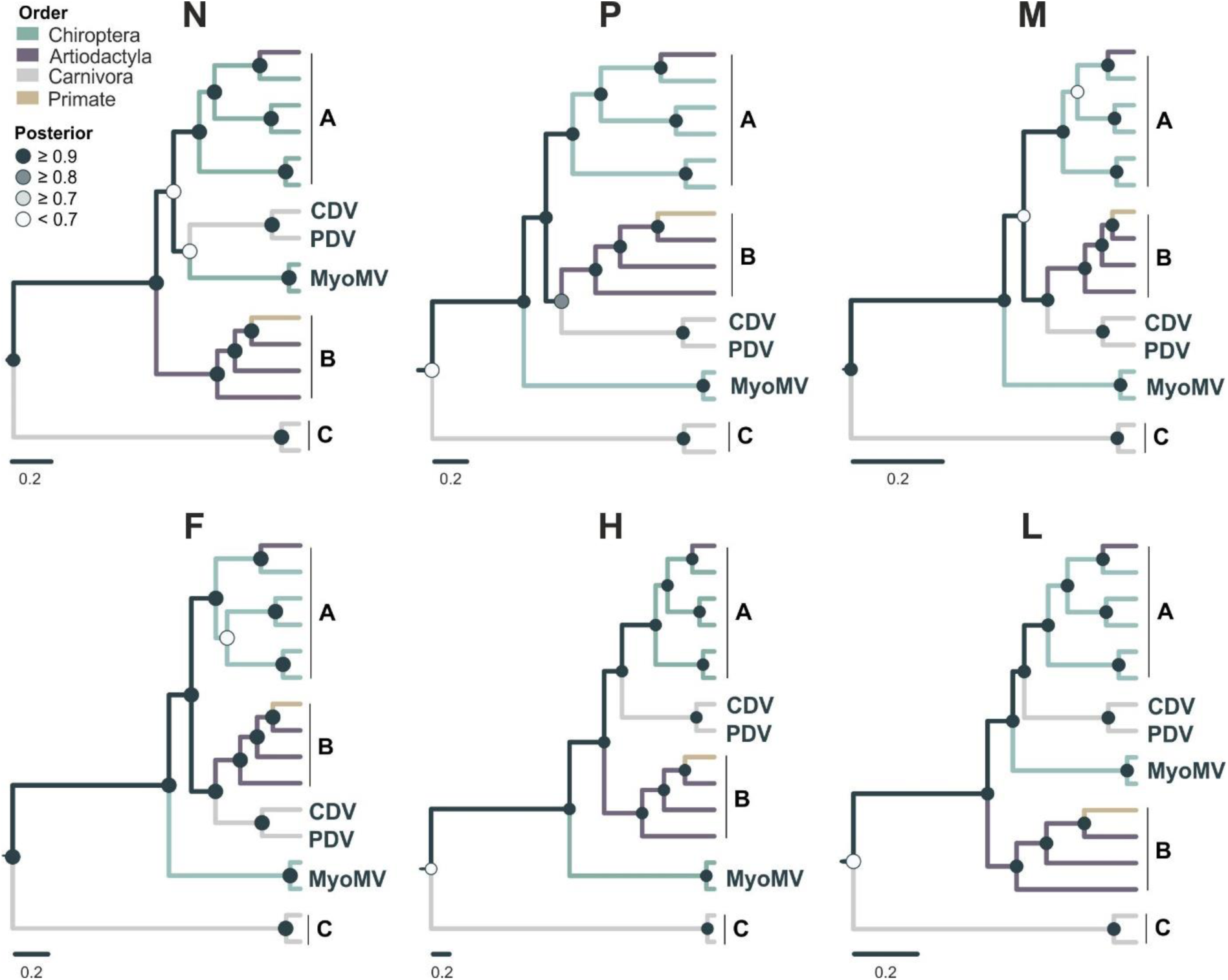
Bayesian phylogenies of individual morbillivirus open reading frames. Trees were generated using translated open reading frames. Bayesian posterior probabilities ≥0.9 are shown as circles at nodes. Scale bar indicates genetic distance. The outgroup used was Longquan Berylmys bowersi morbillivirus 1 (GenBank accession no. MZ328284). Clades a) PoMV, PhyMV, MolMV-1, MolMV-2, DesMV-1, DesMV-2; b) MV, RPV, PPRV, and CeMV; and c) FeMV-1 and FeMV-2. See **Supplementary Tables S1** and **S2** for virus abbreviations and GenBank accession nos.

**Extended Data Fig. 3.**
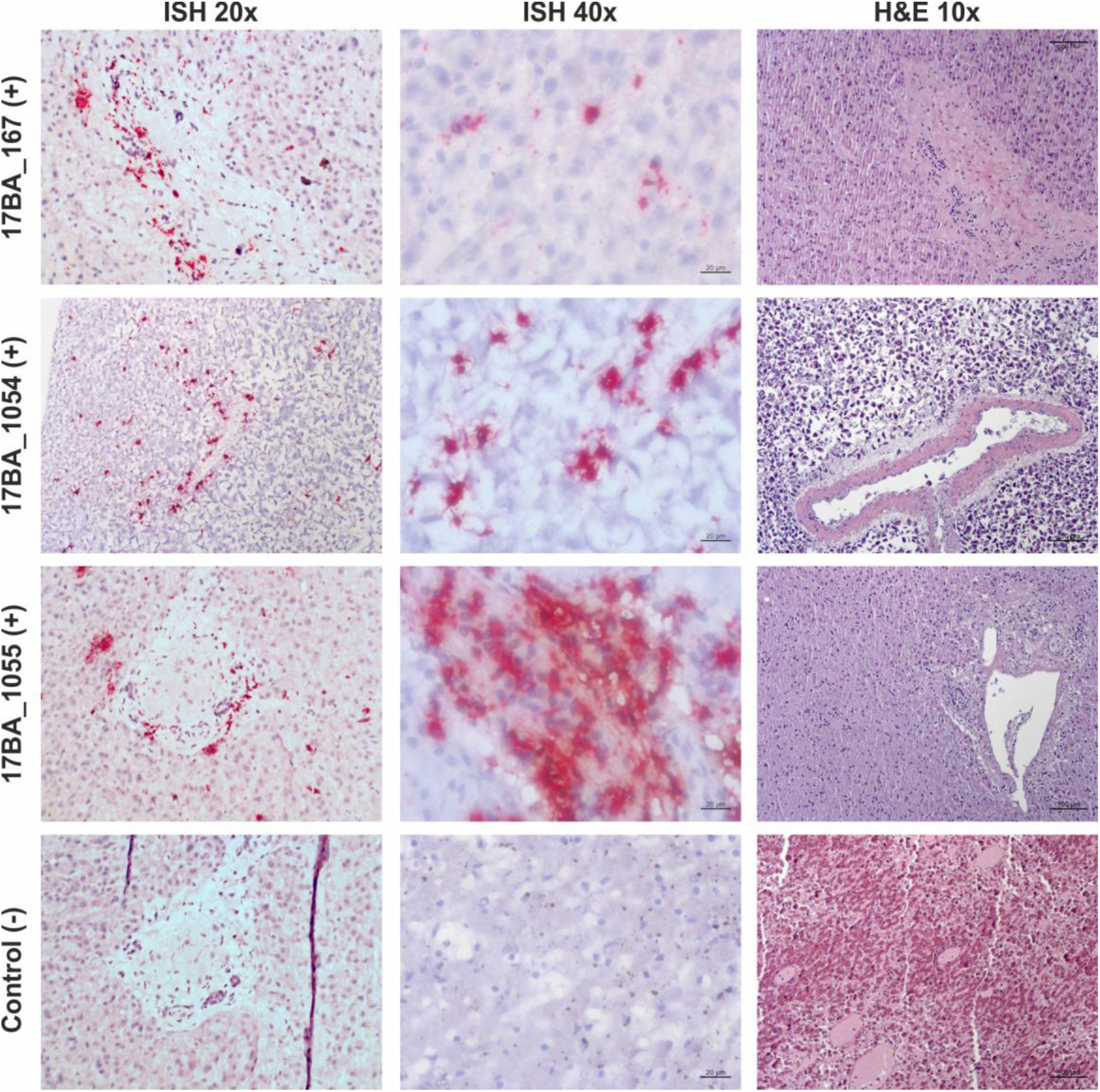
*In situ* hybridization. Strong multifocal MarMV RNA signal (red staining) is shown throughout the periportal space with endothelial portal vein distribution, and intracytoplasmic signal in the epithelial bile duct and hepatocytes in marmosets 17BA_1051, 17BA_1054, and 17BA_1055. No significant histopathological findings were observed in hematoxylin and eosin (H&E) section. No staining was observed in the MarMV-negative animal used as control. (+), MarMV-positive animal, (-), MarMV-negative animal.

**Extended Data Fig. 4.**
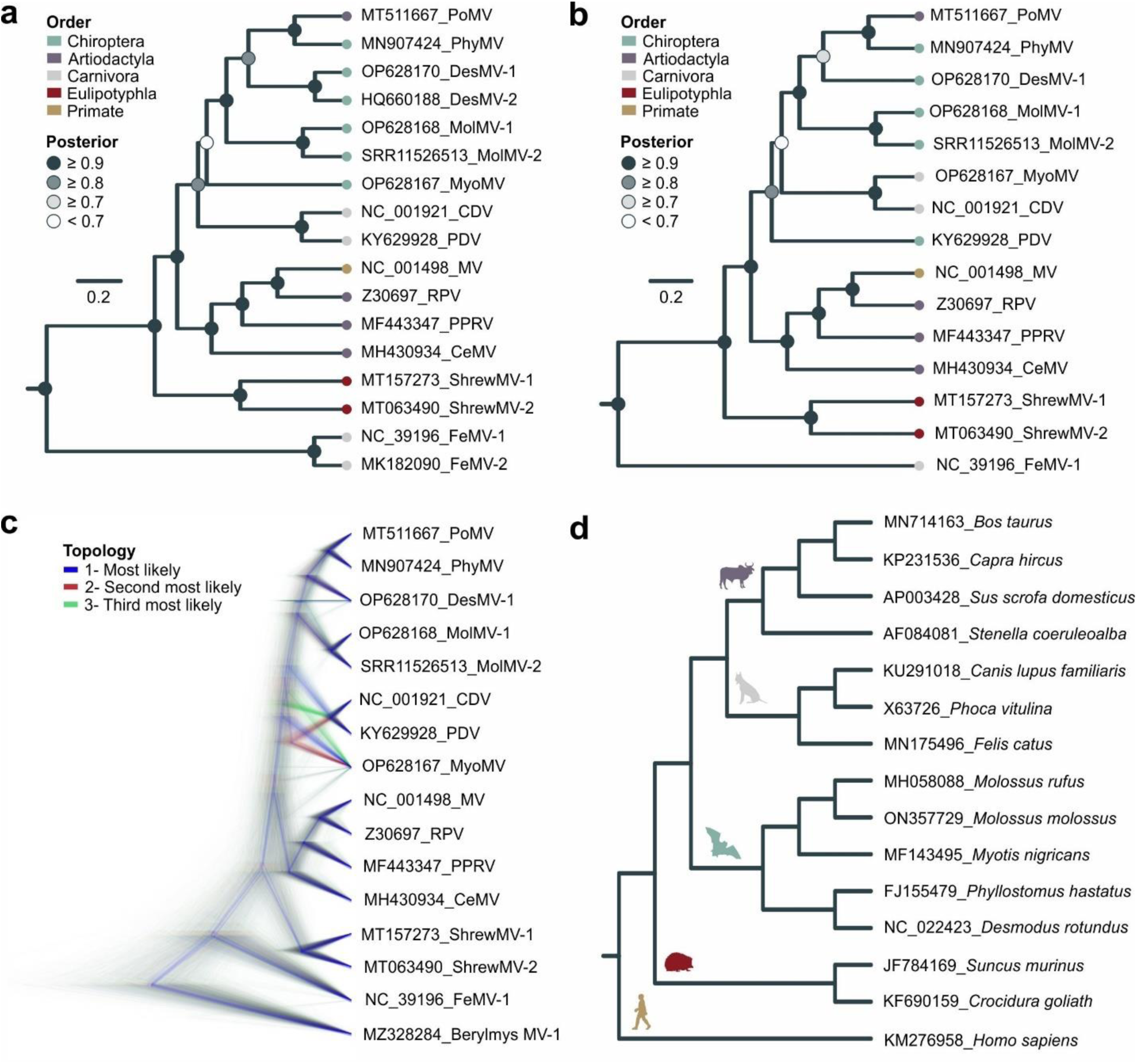
Bayesian phylogeny of morbilliviruses used for coevolutionary analyses. a) Bayesian consensus tree of morbillivirus species using partial *L* gene (528 nucleotides). b) Bayesian consensus tree of morbillivirus species (one virus per host species) using partial *L* gene (528 nucleotides). c) DensiTree showing clustering morbillivirus sequences used to generate the three most likely topologies for event-based cophylogenies. d) Cladogram of morbillivirus host species. See **Supplementary Tables S1-S2 and S4** for virus abbreviations.

**Extended Data Fig. 5.**
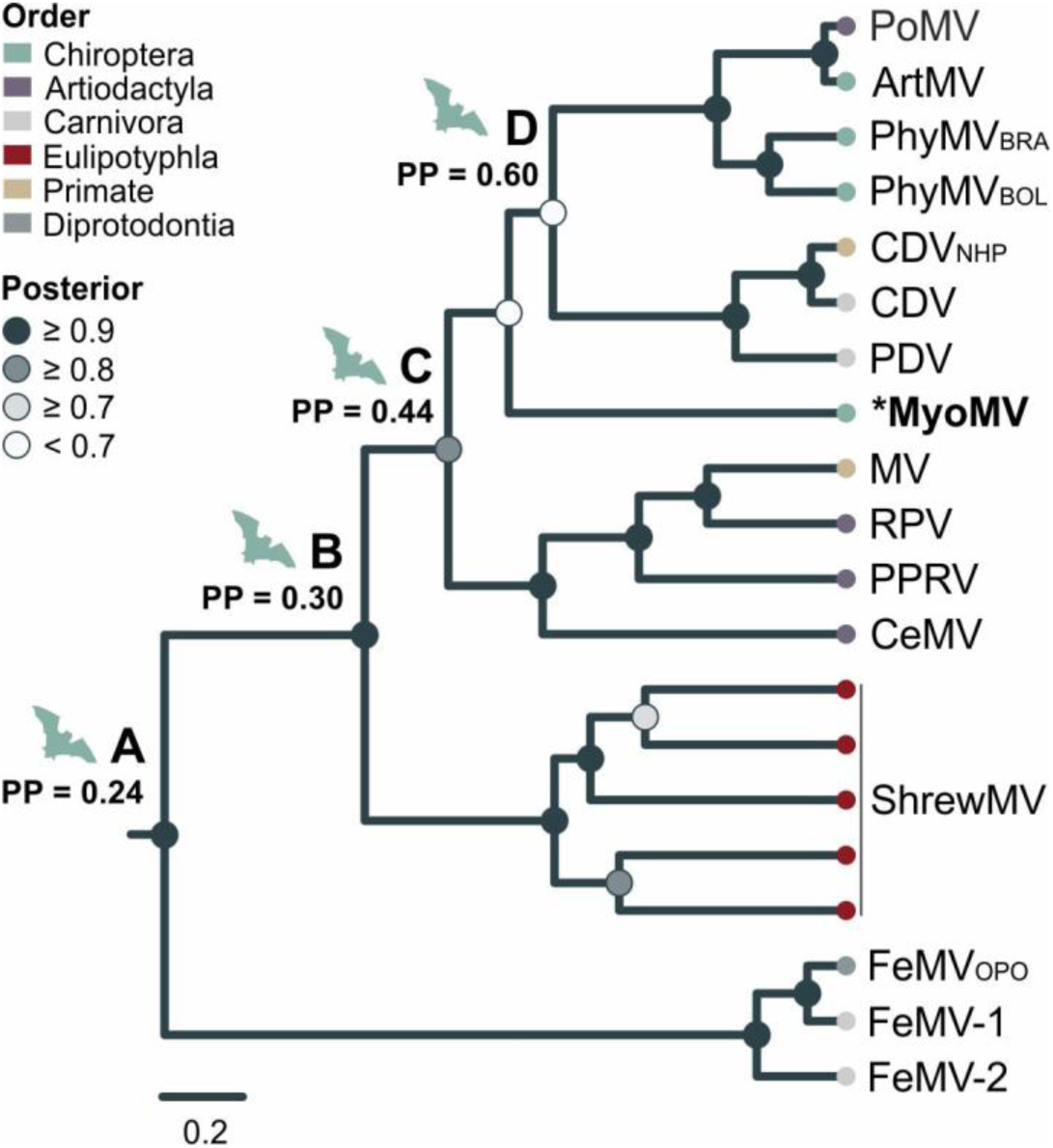
Bayesian phylogeny of previously published classical and divergent morbillivirus species and sequences from suspected cross-order spillover events. Tree was based on partial *L* gene (361 nucleotides). Scale bar indicates genetic distance. At nodes A-D, posterior probabilities (PP) for a bat origin by Bayesian ancestral state reconstructions. BOL, Bolivia; BRA, Brazil; NHP, non-human primate; OPO, opposum. See **Supplementary Table S1** for virus abbreviations and GenBank accession nos.

**Extended Data Fig. 6.**
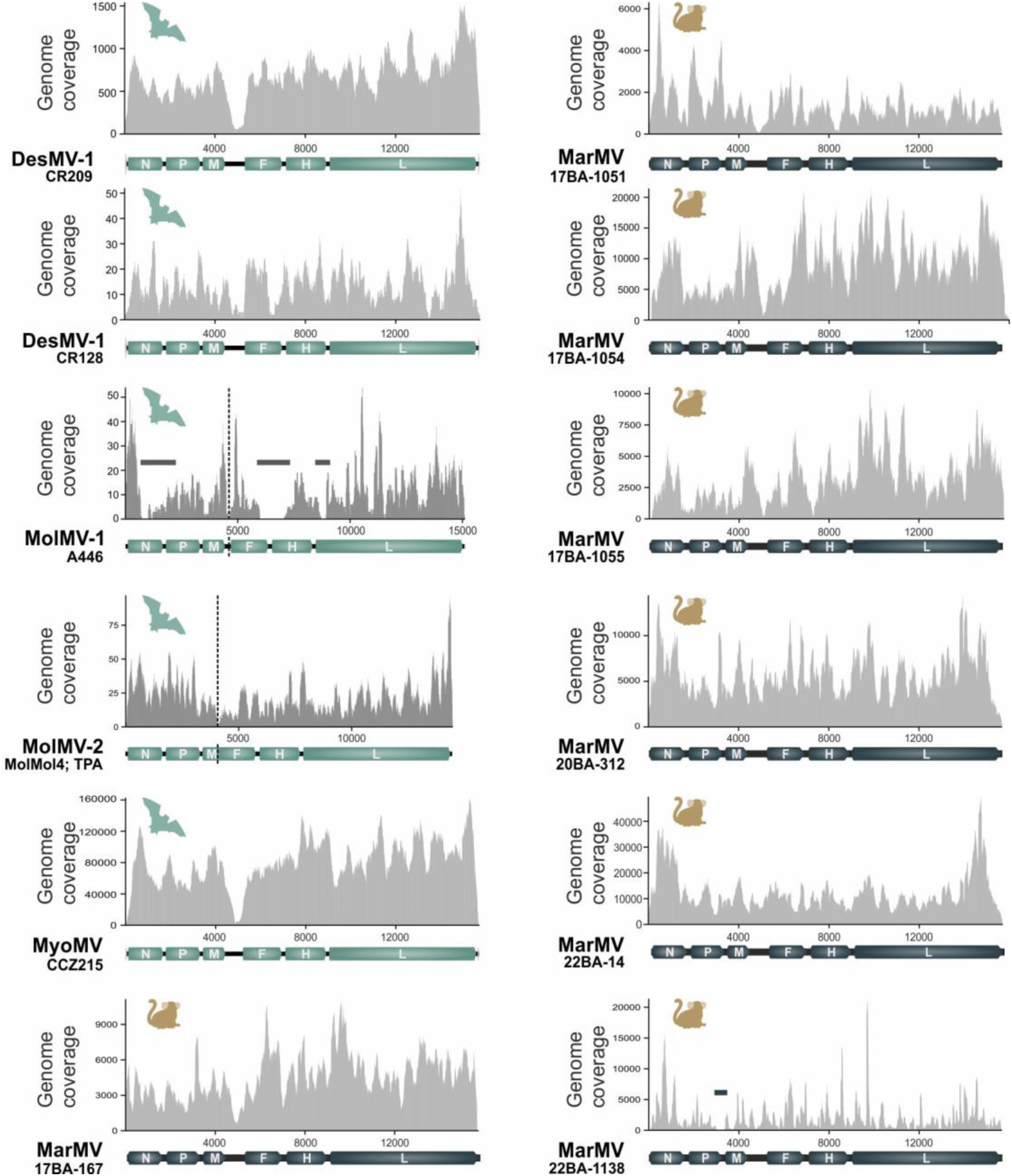
Genome coverage. Genome coverage of assembled genomes and their mean coverage values. Dashed line represents gaps and horizontal line represents genomic regions recovered by Sanger sequencing.

**Extended Data Table 1.**
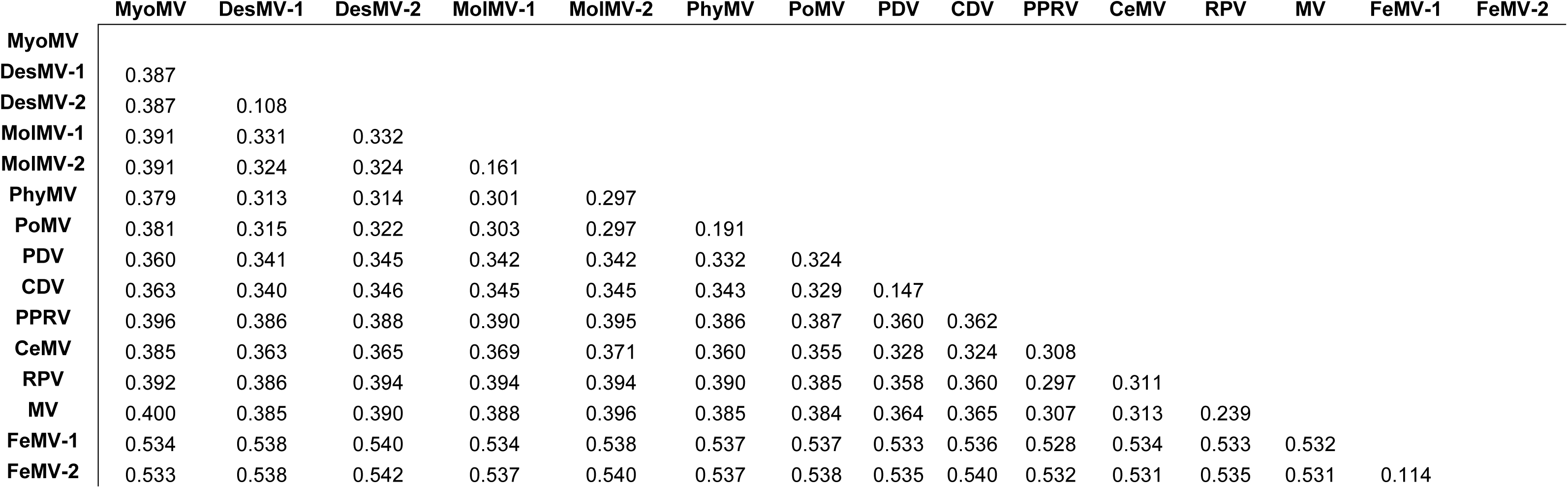
Pairwise amino acid sequence distances of morbilliviruses based on translated, concatenated open reading frames.

|  | MyoMV | DesMV-1 | DesMV-2 | MolMV-1 | MolMV-2 | PhyMV | PoMV | PDV | CDV | PPRV | CeMV | RPV | MV | FeMV-1 | FeMV-2 |
| --- | --- | --- | --- | --- | --- | --- | --- | --- | --- | --- | --- | --- | --- | --- | --- |
| **MyoMV** |  |  |  |  |  |  |  |  |  |  |  |  |  |  |  |
| **DesMV-1** | 0.387 |  |  |  |  |  |  |  |  |  |  |  |  |  |  |
| **DesMV-2** | 0.387 | 0.108 |  |  |  |  |  |  |  |  |  |  |  |  |  |
| **MolMV-1** | 0.391 | 0.331 | 0.332 |  |  |  |  |  |  |  |  |  |  |  |  |
| **MolMV-2** | 0.391 | 0.324 | 0.324 | 0.161 |  |  |  |  |  |  |  |  |  |  |  |
| **PhyMV** | 0.379 | 0.313 | 0.314 | 0.301 | 0.297 |  |  |  |  |  |  |  |  |  |  |
| **PoMV** | 0.381 | 0.315 | 0.322 | 0.303 | 0.297 | 0.191 |  |  |  |  |  |  |  |  |  |
| **PDV** | 0.360 | 0.341 | 0.345 | 0.342 | 0.342 | 0.332 | 0.324 |  |  |  |  |  |  |  |  |
| **CDV** | 0.363 | 0.340 | 0.346 | 0.345 | 0.345 | 0.343 | 0.329 | 0.147 |  |  |  |  |  |  |  |
| **PPRV** | 0.396 | 0.386 | 0.388 | 0.390 | 0.395 | 0.386 | 0.387 | 0.360 | 0.362 |  |  |  |  |  |  |
| **CeMV** | 0.385 | 0.363 | 0.365 | 0.369 | 0.371 | 0.360 | 0.355 | 0.328 | 0.324 | 0.308 |  |  |  |  |  |
| **RPV** | 0.392 | 0.386 | 0.394 | 0.394 | 0.394 | 0.390 | 0.385 | 0.358 | 0.360 | 0.297 | 0.311 |  |  |  |  |
| **MV** | 0.400 | 0.385 | 0.390 | 0.388 | 0.396 | 0.385 | 0.384 | 0.364 | 0.365 | 0.307 | 0.313 | 0.239 |  |  |  |
| **FeMV-1** | 0.534 | 0.538 | 0.540 | 0.534 | 0.538 | 0.537 | 0.537 | 0.533 | 0.536 | 0.528 | 0.534 | 0.533 | 0.532 |  |  |
| **FeMV-2** | 0.533 | 0.538 | 0.542 | 0.537 | 0.540 | 0.537 | 0.538 | 0.535 | 0.540 | 0.532 | 0.531 | 0.535 | 0.531 | 0.114 |  |

**Extended Data Table 2.** Pairwise nucleotide sequence distances of PhyMV and MarMV complete genomes.

|  | Clade<br>PhyMV | MG | BA-1 |  |  |  | BA-2 |  |
| --- | --- | --- | --- | --- | --- | --- | --- | --- |
|  |  | 10875 | 767 | 2707 | 1593 | 2227 | 4464 | 4910 |
| **PhyMV** |  |  |  |  |  |  |  |  |
| **10875** | 0.0202 |  |  |  |  |  |  |  |
| **767** | 0.0448 | 0.0456 |  |  |  |  |  |  |
| **2707** | 0.0449 | 0.0458 | 0.0005 |  |  |  |  |  |
| **1593** | 0.0456 | 0.0465 | 0.0013 | 0.0015 |  |  |  |  |
| **2227** | 0.0466 | 0.0475 | 0.0026 | 0.0025 | 0.0035 |  |  |  |
| **4464** | 0.0478 | 0.0481 | 0.0194 | 0.0196 | 0.0203 | 0.0215 |  |  |
| **4910** | 0.0478 | 0.0481 | 0.0194 | 0.0196 | 0.0203 | 0.0215 | 0.0003 |  |

PhyMV: GenBank accession no. MZ312422

**Extended Data Table 3.** Ancestral state reconstructions at the root of morbillivirus evolutionary tree by BayesTraits.

| Model | Root ancestor | Log-marginal<br>likelihood | 2Δ Log-marginal<br>likelihood | Log(BF) |
| --- | --- | --- | --- | --- |
| M0 (unrestricted) | Unrestricted | -5.128084 |  |  |
| M1 (Fossilised) | Euarchontoglires | -8.396262 | M0-M1 | 6.536356 |
| M2 (Fossilised) | Laurasiatheria | -5.167307 | M0-M2 | 0.078446 |
|  | Euarchontoglires vs. Laurasiatheria |  | M1-M2 | -6.45791 |
|  | Laurasiatheria vs. Euarchontoglires |  | M2-M1 | 6.45791 |

BF, Bayes Factor

**Extended Data Table 4.** Geographic association of bat-, marmoset-, and swine-associated morbilliviruses by BaTS.

| Statistic / MC<br>traits (location) | null mean<br>(95% HDP CIs) | p value |
| --- | --- | --- |
| **AI** | 0.62 (0.3, 0.8) | **0.00** |
| **PS** | 2.95 (2.8, 3.0) | **0.01** |
| **MC (Brazil)** | 5.51 (3.1, 10.0) | **0.01** |
| **MC (Bolivia)** | 1.00 (1.0, 1.0) | 1.00 |
| **MC (Mexico)** | 1.05 (1.0, 1.2) | **0.02** |

MC, monophyletic clade; HPD CIs = highest posterior density confidence intervals (credible sets)

## Notes

### Competing Interest Statement

The authors have declared no competing interest.

### Summary of Updates

This version of the manuscript has been revised to update the license.

## References

1 Rima, B. et al. ICTV Virus Taxonomy Profile: Paramyxoviridae. J Gen Virol 100, 1593–1594 (2019). 10.1099/jgv.0.001328

2 Pastoret, P.-P. et al. in Rinderpest and Peste des Petits Ruminants (eds Thomas Barrett, Paul-Pierre Pastoret, & William P. Taylor) 86–VI (Academic Press, 2006).

3 WHO. Immunization Agenda 2030: A Global Strategy To Leave No One Behind. 60 (2020).

4. WHO. Meales, <https://www.who.int/news-room/fact-sheets/detail/measles> (2019).

5. OIE & FAO. Global Strategy for the Control and Eradication of PPR, <http://www.fao.org/3/a-i4460e.pdf> (2015).

6 Viana, M. et al. Dynamics of a morbillivirus at the domestic-wildlife interface: Canine distemper virus in domestic dogs and lions. Proc Natl Acad Sci U S A 112, 1464–1469 (2015). 10.1073/pnas.1411623112

7 Nambulli, S. et al. FeMV is a cathepsin-dependent unique morbillivirus infecting the kidneys of domestic cats. Proc Natl Acad Sci U S A 119, e2209405119 (2022). 10.1073/pnas.2209405119

8 Arruda, B., Shen, H., Zheng, Y. & Li, G. Novel Morbillivirus as Putative Cause of Fetal Death and Encephalitis among Swine. Emerg Infect Dis 27, 1858–1866 (2021). 10.3201/eid2707.203971

9 Wells, H. L. et al. Classification of new morbillivirus and jeilongvirus sequences from bats sampled in Brazil and Malaysia. Arch Virol (2022). 10.1007/s00705-022-05500-z

10 Drexler, J. F. et al. Bats host major mammalian paramyxoviruses. Nat Commun 3, 796 (2012). 10.1038/ncomms1796

11 Loh, E. H. et al. Prevalence of bat viruses associated with land-use change in the Atlantic Forest, Brazil. Front Cell Infect Microbiol 12, 921950 (2022). 10.3389/fcimb.2022.921950

12 Sasaki, M. et al. Molecular epidemiology of paramyxoviruses in Zambian wild rodents and shrews. J Gen Virol 95, 325–330 (2014). 10.1099/vir.0.058404-0

13 Pinheiro, L. R. S. et al. Identification of Viruses in Molossus Bats from the Brazilian Amazon: A Descriptive Metagenomic Analysis. Microorganisms 12 (2024). 10.3390/microorganisms12030593

14 Alvarez-Carretero, S. et al. A species-level timeline of mammal evolution integrating phylogenomic data. Nature 602, 263–267 (2022). 10.1038/s41586-021-04341-1

15 Irving, A. T., Ahn, M., Goh, G., Anderson, D. E. & Wang, L. F. Lessons from the host defences of bats, a unique viral reservoir. Nature 589, 363–370 (2021). 10.1038/s41586-020-03128-0

16 de Vries, R. D. et al. Delineating morbillivirus entry, dissemination and airborne transmission by studying in vivo competition of multicolor canine distemper viruses in ferrets. PLoS Pathog 13, e1006371 (2017). 10.1371/journal.ppat.1006371

17 Edgar, R. C. et al. Petabase-scale sequence alignment catalyses viral discovery. Nature 602, 142–147 (2022). 10.1038/s41586-021-04332-2

18 Drexler, J. F., Corman, V. M. & Drosten, C. Ecology, evolution and classification of bat coronaviruses in the aftermath of SARS. Antiviral Res 101, 45–56 (2014). 10.1016/j.antiviral.2013.10.013

19 Puryear, W. et al. Longitudinal analysis of pinnipeds in the northwest Atlantic provides insights on endemic circulation of phocine distemper virus. Proc Biol Sci 288, 20211841 (2021). 10.1098/rspb.2021.1841

20 Drexler, J. F. et al. Amplification of emerging viruses in a bat colony. Emerg Infect Dis 17, 449–456 (2011). 10.3201/eid1703.100526

21 Plowright, R. K. et al. Ecological dynamics of emerging bat virus spillover. Proc Biol Sci 282, 20142124 (2015). 10.1098/rspb.2014.2124

22 Nambulli, S., Sharp, C. R., Acciardo, A. S., Drexler, J. F. & Duprex, W. P. Mapping the evolutionary trajectories of morbilliviruses: what, where and whither. Curr Opin Virol 16, 95–105 (2016). 10.1016/j.coviro.2016.01.019

23 Milles, S. et al. An ultraweak interaction in the intrinsically disordered replication machinery is essential for measles virus function. Sci Adv 4, eaat7778 (2018). 10.1126/sciadv.aat7778

24 Simon-Loriere, E. & Holmes, E. C. Why do RNA viruses recombine? Nat Rev Microbiol 9, 617–626 (2011). 10.1038/nrmicro2614

25 Hashiguchi, T. et al. Structure of the measles virus hemagglutinin bound to its cellular receptor SLAM. Nat Struct Mol Biol 18, 135–141 (2011). 10.1038/nsmb.1969

26 Ikegame, S. et al. Metagenomics-enabled reverse-genetics assembly and characterization of myotis bat morbillivirus. Nat Microbiol 8, 1108–1122 (2023). 10.1038/s41564-023-01380-4

27 de Vries, R. D. et al. Measles vaccination of nonhuman primates provides partial protection against infection with canine distemper virus. J Virol 88, 4423–4433 (2014). 10.1128/JVI.03676-13

28 de Sousa, L. L. F. et al. Rabies virus variants from bats closely related to variants found in marmosets (Callithrix jacchus), a neglected source of human rabies infection in Brazil. J Med Virol 95, e29046 (2023). 10.1002/jmv.29046

29 Amman, B. R., Schuh, A. J., Albarino, C. G. & Towner, J. S. Marburg Virus Persistence on Fruit as a Plausible Route of Bat to Primate Filovirus Transmission. Viruses 13 (2021). 10.3390/v13122394

30 Mares-Guia, M. et al. Yellow fever epizootics in non-human primates, Southeast and Northeast Brazil (2017 and 2018). Parasit Vectors 13, 90 (2020). 10.1186/s13071-020-3966-x

31 Dinh, A., Fleuret, V. & Hanslik, T. Liver involvement in adults with measles. Int J Infect Dis 17, e1243–1244 (2013). 10.1016/j.ijid.2013.06.014

32 Kadam, R. G. et al. Molecular and pathological screening of canine distemper virus in Asiatic lions, tigers, leopards, snow leopards, clouded leopards, leopard cats, jungle cats, civet cats, fishing cat, and jaguar of different states, India. Infect Genet Evol 98, 105211 (2022). 10.1016/j.meegid.2022.105211

33 Foley, N. M., Springer, M. S. & Teeling, E. C. Mammal madness: is the mammal tree of life not yet resolved? Philos Trans R Soc Lond B Biol Sci 371 (2016). 10.1098/rstb.2015.0140

34 Dux, A. et al. Measles virus and rinderpest virus divergence dated to the sixth century BCE. Science 368, 1367–1370 (2020). 10.1126/science.aba9411

35 Martinez-Gutierrez, M. & Ruiz-Saenz, J. Diversity of susceptible hosts in canine distemper virus infection: a systematic review and data synthesis. BMC Vet Res 12, 78 (2016). 10.1186/s12917-016-0702-z

36 Lavorente, F. L. P. et al. First detection of Feline morbillivirus infection in white-eared opossums (Didelphis albiventris, Lund, 1840), a non-feline host. Transbound Emerg Dis 69, 1426–1437 (2022). 10.1111/tbed.14109

37 Jo, W. K. et al. Evolutionary evidence for multi-host transmission of cetacean morbillivirus. Emerg Microbes Infect 7, 201 (2018). 10.1038/s41426-018-0207-x

38 Olival, K. J. et al. Host and viral traits predict zoonotic spillover from mammals. Nature 546, 646–650 (2017). 10.1038/nature22975

39 Lau, S. K. et al. Identification and complete genome analysis of three novel paramyxoviruses, Tuhoko virus 1, 2 and 3, in fruit bats from China. Virology 404, 106–116 (2010). 10.1016/j.virol.2010.03.049

40 Rojas, D., Warsi, O. M. & Davalos, L. M. Bats (Chiroptera: Noctilionoidea) Challenge a Recent Origin of Extant Neotropical Diversity. Syst Biol 65, 432–448 (2016). 10.1093/sysbio/syw011

41 Lack, J. B., Roehrs, Z. P., Stanley, C. E., M., R. & van den Bussche, R. A. Molecular phylogenetics of Myotis indicate familial-level divergence for the genus Cistugo (Chiroptera). Journal of Mammalogy 91, 976–992 (2010). 10.1644/09-MAMM-A-192.1

42 Ammerman, L. K., Lee, D. N. & Tipps, T. M. First molecular phylogenetic insights into the evolution of free-tailed bats in the subfamily Molossinae (Molossidae, Chiroptera). Journal of Mammalogy 93, 12–28 (2012). 10.1644/11-mamm-a-103.1

43. Dias Sousa, A. V. Beefing up: meeting Top 10 Beef Producing Countries Worldwide, <https://ruminants.ceva.pro/beef-producing-countries> (2024).

44 Shahbandeh, M. Number of pigs worldwide in 2023, by country, <https://www.statista.com/statistics/263964/number-of-pigs-in-selected-countries/#:~:text=Global%20overview,tons%20of%20pork%20each%20year.> (2023).

45 Field, H. et al. The natural history of Hendra and Nipah viruses. Microbes Infect 3, 307–314 (2001). 10.1016/s1286-4579(01)01384-3

46 Zhou, P. et al. Fatal swine acute diarrhoea syndrome caused by an HKU2-related coronavirus of bat origin. Nature 556, 255–258 (2018). 10.1038/s41586-018-0010-9

47 Hernandez-Mora, G. et al. Virulent Brucella nosferati infecting Desmodus rotundus has emerging potential due to the broad foraging range of its bat host for humans and wild and domestic animals. mSphere 8, e0006123 (2023). 10.1128/msphere.00061-23

48 Schneider, M. C. et al. Rabies transmitted by vampire bats to humans: an emerging zoonotic disease in Latin America? Rev Panam Salud Publica 25, 260–269 (2009). 10.1590/s1020-49892009000300010

49 Gurley, E. S. et al. Person-to-person transmission of Nipah virus in a Bangladeshi community. Emerg Infect Dis 13, 1031–1037 (2007). 10.3201/eid1307.061128

50 Lizarro, D., Galarza, M. I. & Aguirre, L. F. Traffic and trade of Bolivian bats. Rev. Bol. Ecol. y Cons. Amb. 27, 63–75 (2010).

51 FAO & OIE. Global Rinderpest Action Plan - Post-Eradication, 2018).

52 WOAH. Extension to the Designation of Facilities Holding Rinderpest Virus Containing Material to Maintain Global Freedom from Rinderpest. 1–5 (2022).

53 Holzer, B., Hodgson, S., Logan, N., Willett, B. & Baron, M. D. Protection of Cattle against Rinderpest by Vaccination with Wild-Type but Not Attenuated Strains of Peste des Petits Ruminants Virus. J Virol 90, 5152–5162 (2016). 10.1128/JVI.00040-16

54 Schulz, C., Fast, C., Schlottau, K., Hoffmann, B. & Beer, M. Neglected Hosts of Small Ruminant Morbillivirus. Emerg Infect Dis 24, 2334–2337 (2018). 10.3201/eid2412.180507

55 Tan, C. W., Valkenburg, S. A., Poon, L. L. M. & Wang, L. F. Broad-spectrum pan-genus and pan-family virus vaccines. Cell Host Microbe 31, 902–916 (2023). 10.1016/j.chom.2023.05.017

56 Isidro, J. et al. Phylogenomic characterization and signs of microevolution in the 2022 multi-country outbreak of monkeypox virus. Nat Med (2022). 10.1038/s41591-022-01907-y

57 Alcaide, M. et al. Disentangling vector-borne transmission networks: a universal DNA barcoding method to identify vertebrate hosts from arthropod bloodmeals. PLoS One 4, e7092 (2009). 10.1371/journal.pone.0007092

58 Langmead, B. & Salzberg, S. L. Fast gapped-read alignment with Bowtie 2. Nat Methods 9, 357–359 (2012). 10.1038/nmeth.1923

59 Tong, S., Chern, S. W., Li, Y., Pallansch, M. A. & Anderson, L. J. Sensitive and broadly reactive reverse transcription-PCR assays to detect novel paramyxoviruses. J Clin Microbiol 46, 2652–2658 (2008). 10.1128/JCM.00192-08

60 Seki, F. & Takeda, M. Novel and classical morbilliviruses: Current knowledge of three divergent morbillivirus groups. Microbiol Immunol 66, 552–563 (2022). 10.1111/1348-0421.13030

61 Tatsuo, H., Ono, N. & Yanagi, Y. Morbilliviruses use signaling lymphocyte activation molecules (CD150) as cellular receptors. J Virol 75, 5842–5850 (2001). 10.1128/JVI.75.13.5842-5850.2001

62 Martin, D. P., Murrell, B., Golden, M., Khoosal, A. & Muhire, B. RDP4: Detection and analysis of recombination patterns in virus genomes. Virus Evol 1, vev003 (2015). 10.1093/ve/vev003

63. Santichaivekin, S., et al. eMPRess: a systematic cophylogeny reconciliation tool. Bioinformatics 37, 2481–2482 (2021). 10.1093/bioinformatics/btaa978

64 Bouckaert, R. R. DensiTree: making sense of sets of phylogenetic trees. Bioinformatics 26, 1372–1373 (2010). 10.1093/bioinformatics/btq110

65 Parker, J., Rambaut, A. & Pybus, O. G. Correlating viral phenotypes with phylogeny: accounting for phylogenetic uncertainty. Infect Genet Evol 8, 239–246 (2008). 10.1016/j.meegid.2007.08.001

66 Meade, A. & Pagel, M. Ancestral State Reconstruction Using BayesTraits. Methods Mol Biol 2569, 255–266 (2022). 10.1007/978-1-0716-2691-7_12

67 Goes, L. G. et al. Novel bat coronaviruses, Brazil and Mexico. Emerg Infect Dis 19, 1711–1713 (2013). 10.3201/eid1910.130525

68 Goes, L. G. B. et al. Genetic diversity of bats coronaviruses in the Atlantic Forest hotspot biome, Brazil. Infect Genet Evol 44, 510–513 (2016). 10.1016/j.meegid.2016.07.034

69 Campos, A. C. A. et al. Bat Influenza A(HL18NL11) Virus in Fruit Bats, Brazil. Emerg Infect Dis 25, 333–337 (2019). 10.3201/eid2502.181246

70 Carneiro, A. J. et al. Rabies virus RNA in naturally infected vampire bats, northeastern Brazil. Emerg Infect Dis 16, 2004–2006 (2010). 10.3201/eid1612.100726

71 Mioni, M. S. R. et al. Septicemia due to Streptococcus dysgalactiae subspecies dysgalactiae in vampire bats (Desmodus rotundus). Sci Rep 8, 9772 (2018). 10.1038/s41598-018-28061-1

